# Neural circuit-wide analysis of gene expression during deafening-induced destabilization of birdsong

**DOI:** 10.1101/2022.12.13.520194

**Authors:** Bradley M. Colquitt, Kelly Li, Foad Green, Robert Veline, Michael S. Brainard

**Author notes:** Current Affiliation: Department of Molecular, Cell, and Developmental Biology, University of California-Santa Cruz, Santa Cruz, CA, 95064. Current Affiliation: Syapse, Inc., San Francisco, CA, 94115. Current Affiliation: The Advanced Science Research Center, The City University of New York; The Graduate Center at the City University of New York, New York, NY, 10016. Corresponding authors: Bradley M. Colquitt, Department of Molecular, Cell, and Developmental Biology, University of California-Santa Cruz 1156 High St., Sinsheimer Labs 329 Santa Cruz, CA 95064; Michael S. Brainard, Howard Hughes Medical Institute and Department of Physiology, University of California-San Francisco, 675 Nelson Rising Ln Room 540, Box 0444 San Francisco, CA 94158.

## Abstract

Sensory feedback is required for stable execution of learned motor skills, and its loss can severely disrupt motor performance. The neural mechanisms that mediate sensorimotor stability have been extensively studied at systems and physiological levels, yet relatively little is known about how disruptions to sensory input alter the molecular properties of associated motor systems. Songbird courtship song, a model for skilled behavior, is a learned and highly structured vocalization that is destabilized following deafening. Here, we sought to determine how the loss of auditory feedback modifies gene expression and its coordination across the birdsong sensorimotor circuit. To facilitate this system-wide analysis of transcriptional responses, we developed a gene expression profiling approach that enables the construction of hundreds of spatially-defined RNA-sequencing libraries. Using this method, we found that deafening preferentially alters gene expression across birdsong neural circuitry relative to surrounding areas, particularly in premotor and striatal regions. Genes with altered expression are associated with synaptic transmission, neuronal spines, and neuromodulation and show a bias toward expression in glutamatergic neurons and *Pvalb/Sst-*class GABAergic interneurons. We also found that connected song regions exhibit correlations in gene expression that were reduced in deafened birds relative to hearing birds, suggesting that song destabilization alters the inter-region coordination of transcriptional state. Finally, lesioning LMAN, a forebrain afferent of RA required for deafening-induced song plasticity, had the largest effect on groups of genes that were also most affected by deafening. Combined, this integrated transcriptomics analysis demonstrates that the loss of peripheral sensory input drives a distributed gene expression response throughout associated sensorimotor neural circuitry and identifies specific candidate molecular and cellular mechanisms that support stability and plasticity of learned motor skills.

## Introduction

The accurate and stable performance of motor skills relies on sensory feedback (Todorov, 2004). The loss of this feedback, for example through hearing or vision loss from injury or neurodegeneration, can lead to increased errors in the execution of even well-learned motor behaviors, such as speech and walking (Lane and Webster, 1991; Waldstein, 1990; Wood et al., 2011). Yet it is poorly understood how such peripheral sensory loss influences the properties of central motor circuits to drive neural plasticity and how these effects in turn influence motor output.

The courtship song of songbirds, a learned motor skill subserved by a dedicated and discrete neural architecture, offers a tractable system in which to characterize the neural mechanisms that underlie sensorimotor integration and motor skill stability. Juvenile birds produce unstructured and variable vocalizations that, over the course of several months of learning, become more structured, less variable, and more similar to adult song (Brainard and Doupe, 2013). In finches, after this developmental learning period, birdsong performance remains extraordinarily consistent from rendition-to-rendition over the course of a bird’s life and is said to be ‘crystallized’. However, auditory feedback plays an essential role in maintaining this stability; modifying auditory feedback or completely removing auditory input through deafening, can drive changes to song (Brainard and Doupe, 2001, 2000; Fukushima and Margoliash, 2015; Leonardo and Konishi, 1999; Lombardino and Nottebohm, 2000; Nordeen and Nordeen, 1992; Okanoya and Yamaguchi, 1997; Tschida and Mooney, 2012; Woolley and Rubel, 1997). The neural mechanisms that underlie these changes have been studied in terms of physiology and morphology, yet we lack a transcriptome- and circuit-wide understanding of how altered auditory feedback influences gene expression in song sensorimotor circuitry — a critical biological vantage point to understand how sensory information intersects with nervous system to influence motor plasticity.

To gain insight into the molecular and cellular factors that regulate the stability of adult birdsong, we analyzed gene expression alterations in birdsong sensorimotor circuitry and surrounding non-song regions in response to deafening, a strong driver of song destabilization. We developed a gene expression profiling approach that enabled large-scale analysis of gene expression in spatially defined brain regions. Using this technique, we identified a suite of expression changes across the song system, including region-specific and common transcriptional responses as well as altered gene expression correlations across regions. Using a previously generated single-cell atlas of the premotor portion of song neural circuitry, we identified the cellular types that experience the greatest transcriptional change following deafening. Finally, we examined how input from a song region required for deafening-induced song plasticity influences gene expression in its song premotor target and found a diverse set of expression changes, with substantial overlap with those elicited by deafening.

## Results

### Deafening destabilizes birdsong and increases song variability

Songbirds rely on auditory feedback to maintain the quality of their songs (Fig. 1A). Past work has shown that experimentally removing this feedback by deafening results in the gradual deterioration of both song spectral structure and temporal ordering of the individual elements (syllables) that comprise song (Nordeen and Nordeen, 1992; Okanoya and Yamaguchi, 1997; Woolley and Rubel, 1997). Deafening also drives a range of physiological, cellular, and molecular changes in the song system, including alterations to neuronal turnover (Scott et al., 2000; Wang et al., 1999) (but see (Pytte et al., 2012)), dendritic spine morphology (Peng et al., 2013, 2012a; Tschida and Mooney, 2012; Zhou et al., 2017), neuronal excitability (Tschida and Mooney, 2012), and gene expression (Watanabe et al., 2002).

**Figure 1.**
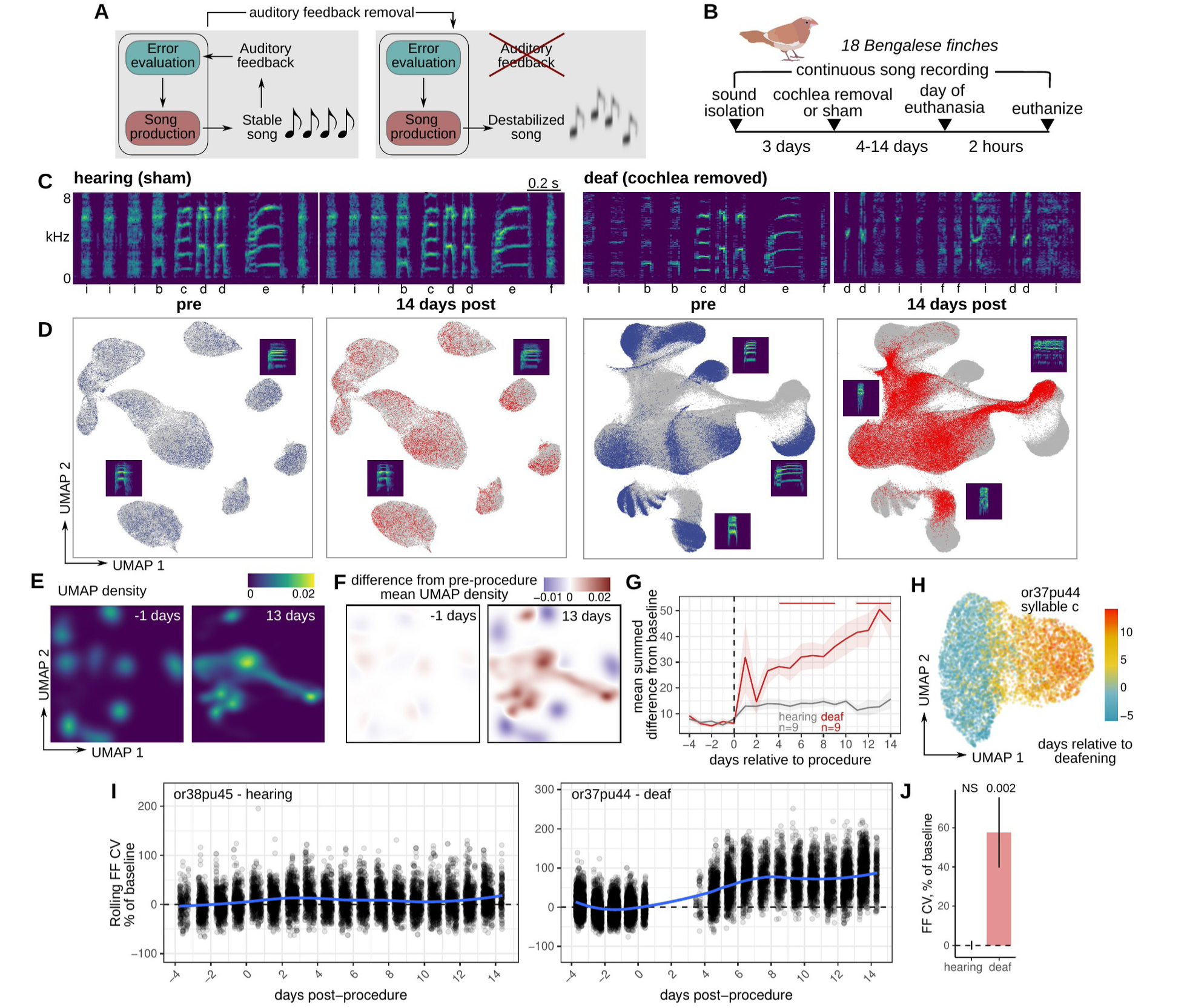
Rapid and global destabilization of song following deafening. **(A)** Song destabilization through the removal of auditory feedback. Adult songbirds use auditory feedback to evaluate their own song production and maintain song quality and consistency. Loss of auditory feedback results in the gradual destabilization of song. **(B)** Experimental overview. After a baseline period of song recording, Bengalese finches (*Lonchura striata domestica*) were either deafened through bilateral cochlea removal or underwent a sham surgery. After 4, 9, or 14 days post-surgery, birds were euthanized for gene expression analysis. Bengalese finch graphic obtained from (Sainburg, 2020). **(C)** Example spectrograms from one hearing (sham) and one deaf (bilateral cochlea removal) bird. Songs are shown from before the procedure and 14 days following the procedure. Labels below each spectrogram correspond to discrete categories of song units (‘syllables’). kHz, kiloHertz. **(D)** Uniform Manifold Approximation and Projection (UMAP) representation of syllable spectrograms (see Methods) across the entire recording period for each bird (4 days before to 14 days after the procedure). Data are split into ‘pre’-procedure (4 to 1 day before surgery) and ‘post’-procedure (1 to 14 days after surgery) subset. For reference, gray points in each plot correspond to data from the other subset. Example syllable spectrograms are placed adjacent to their position in UMAP space. **(E)** Density plots of UMAP projections for the syllables from one deafened bird (shown in panel (A)) at two timepoints, one day before and 13 days after deafening. **(F)** Subtraction of UMAP densities in (E) from the average pre-procedure density. **(G)** Mean sum of UMAP density differences for syllables from birds that were either deafened (deaf) or underwent a sham surgery (hearing). For each bird and each day, positive UMAP density differences were summed then averaged across birds. Error bands are standard errors of the mean. Color bars indicate days in which values were significantly different between deaf and hearing birds (Student’s t-test, two-sided, p<0.05). **(H)** UMAP plot of one syllable from one deafened bird colored by day following deafening. **(I)** Comparison of fundamental frequency (FF) variability between hearing and deafened birds. Rolling coefficient of variation (CV, window size 11 syllables) was calculated for the fundamental frequencies of each harmonic stack for each bird. Shown are two example syllables from one hearing and one deafened bird, plotted across the number of days relative to the procedure date (sham or cochlea removal). **(J)** Mean FF CV in the 7 to 9 days following sham or cochlea removal normalized to FF CV in the 2 days before procedure. Linear mixed-effects regression (see Methods) was used to estimate the group post vs. pre-procedure FF CV difference for hearing and deaf birds. P-values are obtained from the regression model using Satterthwaite’s degrees of freedom method.

**Supplemental Figure 1-1.**
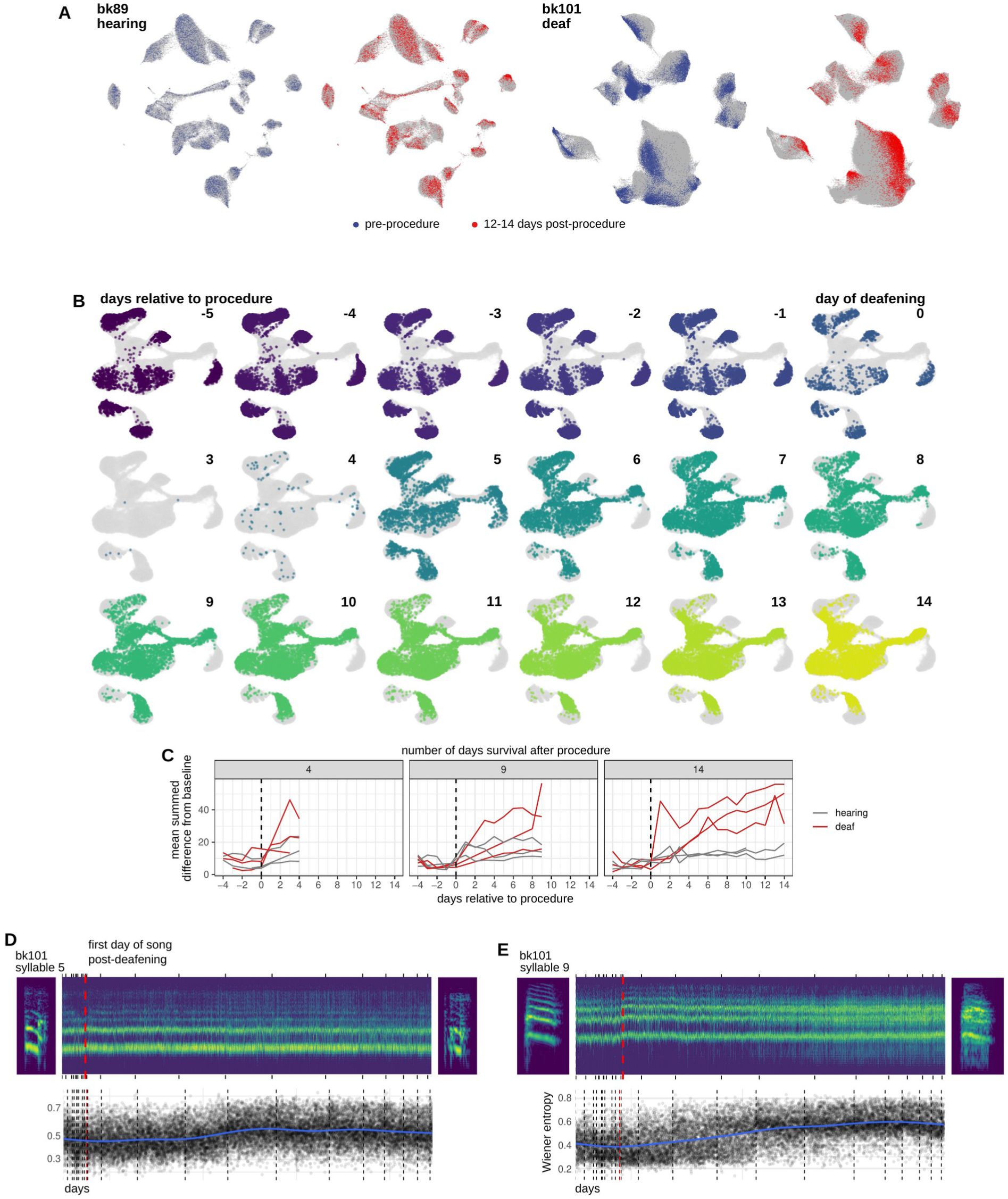
Additional quantification of deafening-induced changes to song. **(A)** UMAP representation of syllable spectrograms (see Methods) across the entire recording period (4 days before to 14 days after the procedure) for an example hearing (bk89) and deafened (bk101) bird. Data are split into ‘pre’-procedure (4 to 1 day before surgery) and ‘post’-procedure (12 to 14 days after surgery) subset. For reference, gray points in each plot correspond to data from other days. Example syllable spectrograms are placed adjacent to their position in UMAP space. **(B)** UMAP representations of syllable spectrograms from a deafened bird split by day relative to cochlea removal. Data used is the same that is presented in Figure 1C,D. **(C)** Break out of the UMAP distance analysis from Figure 1G into individual birds. **(D)** Increase in syllable noise over time following deafening. Example spectrograms for two syllables from the day before (*left*) and 14 days following (*right*) deafening. Spectrograms were averaged along the time axis within red boxed regions shown on the examples and plotted over time. Vertical dashed red line indicates the day of deafening. Tick marks indicate the beginning of each day. Shown is 20% of the total number of syllables. **(E)** Spectral flatness (also known as Wiener entropy) over time. Measure was z-scored relative to pre-procedure. Vertical dashed red line indicates the day of deafening. Vertical dashed black lines indicate the beginning of each day.

We reasoned that comparisons of gene expression in birds undergoing song destabilization following deafening would uncover molecular pathways involved in either promoting or limiting song plasticity. We therefore generated a cohort of eighteen adult male Bengalese finches (*Lonchura striata domestica*) that were either deafened through bilateral cochlea removal or underwent a sham surgery (9 birds each condition, Fig. 1B). This cohort was further divided into sets of birds that were euthanized 4, 9, or 14 days post-procedure (3 birds per procedure type and time point). Birds were euthanized two hours after lights-on. As in previous studies (Brainard and Doupe, 2000; Okanoya and Yamaguchi, 1997; Tschida and Mooney, 2012; Woolley and Rubel, 1997), deafening caused a gradual decay of song quality over the course of several days, while sham surgery induced relatively little song change (Fig. 1B-H, Figure 1-figure supplement 1A-C). To visualize deafening-induced changes to song, we calculated spectrograms for each syllable and used uniform manifold approximate projection (UMAP) to project these spectrograms onto a latent space, using an approach described in Sainburg et al. (Sainburg et al., 2020) (Fig. 1C, Figure 1-figure supplement 1A-C). Following deafening, the projections of syllable spectrograms gradually shifted to occupy different locations, indicating a change from pre-procedure song (Fig. 1D-H, Figure 1-figure supplement 1A,B). Syllable spectral changes after deafening were complex but generally trended toward an increase in syllable ‘noisiness’ (Wiener entropy) (Figure 1-figure supplement 1D,E).

Although adult birdsong is a highly precise motor skill, its features vary slightly from rendition-to-rendition, similar to other motor skills. Past work has demonstrated that this variability is in part generated by central neural mechanisms in the forebrain and is modulated by social context, indicating that song variability is an actively regulated component of birdsong (Kao et al., 2005; Kao and Brainard, 2006; Kojima et al., 2013; Moorman et al., 2021; Sakata et al., 2008). To assess how song variability changes following deafening, we focused on a single spectral feature, fundamental frequency (FF), and calculated its coefficient of variation (CV) across renditions (Fig. 1J,K). Deafening resulted in a gradual increase in the CV of FF (day 7-9 post-procedure mean ± sem, 57 ± 18%) while sham surgery elicited no change (0.11 ± 2.3%). This increase in rendition-to-rendition FF variability is consistent with reports describing an increase in within-syllable frequency modulation following deafening (Brainard and Doupe, 2001). These results indicate that deafening elicits both shifts in the structure of song as well as decreases in stereotypy across renditions.

### Neural circuit wide analysis of gene expression

Birdsong is generated by a dedicated and anatomically discrete neural circuit called the song system (Fig. 2A and B). This defined architecture allows interrogation of how the molecular and cellular properties of each region (termed ‘song nuclei’) influence and are influenced by birdsong performance and learning. Four primary song nuclei reside in the telencephalon: HVC (proper name), RA (robust nucleus of the arcopallium), LMAN (lateral magnocellular nucleus of the anterior nidopallium), and Area X. HVC and RA comprise the song motor pathway (SMP) and are necessary for song performance (Nottebohm et al., 1976; Simpson and Vicario, 1990). HVC influences the timing and temporal structure of song and projects to RA, which provides descending motor control of song via projections to syringeal and respiratory brainstem regions, which send recurrent connections back into the SMP to influence spectral and temporal features of song (Goldberg and Fee, 2012; Vicario, 1991; Wild, 1993). HVC also projects to the striatal nucleus Area X that, together with the pallial region LMAN and thalamic region DLM, form the ’anterior forebrain pathway’ (AFP), which contributes to song plasticity both during song acquisition in juveniles and song adaptation in adults (Andalman and Fee, 2009; Bottjer et al., 1984; Brainard and Doupe, 2000; Charlesworth et al., 2012; Nordeen and Nordeen, 1993; Scharff and Nottebohm, 1991; Sohrabji et al., 1990; Warren et al., 2011; Williams and Mehta, 1999). Each region is embedded in a larger anatomical domain that lies outside of song control circuitry but shares similar molecular, connectivity, and functional properties (Fig. 2B) (Feenders et al., 2008; Helduser et al., 2013; Kröner and Güntürkün, 1999). HVC is located in the dorsal part of the caudal nidopallium (NC); RA is located in the arcopallium (Arco.); Area X is located in the striatum (Stri.); and LMAN is located in the rostral nidopallium (NR). In effect, these regions serve as ‘non-song’ comparators for each song region that enable identification of molecular and cellular features that are specific to song-related perturbations.

**Figure 2.**
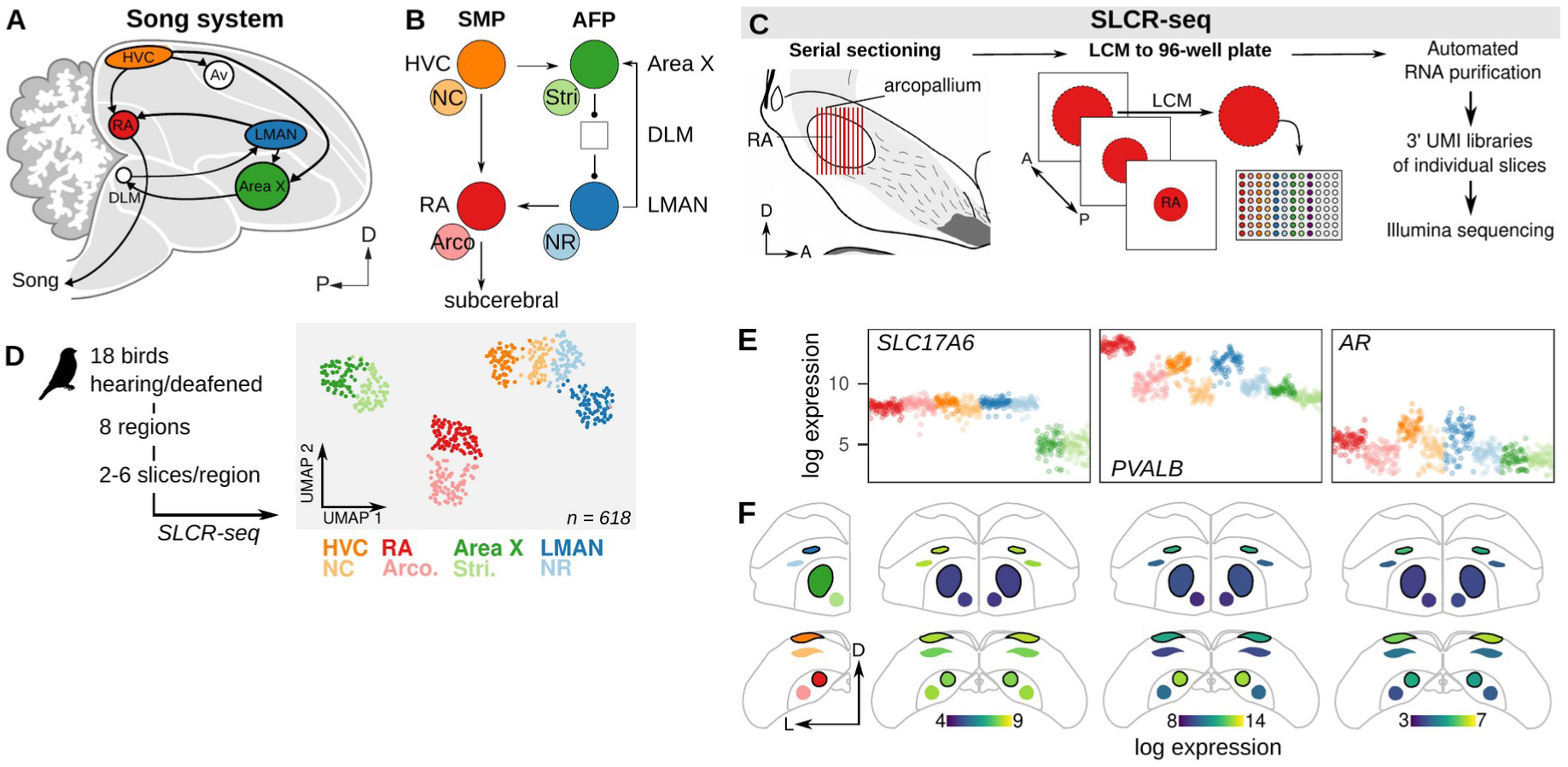
Neural circuit transcriptomics using Serial Laser Capture RNA-sequencing (SLCR-seq). **(A)** Schematic overview of the song system. HVC, proper name; RA, robust nucleus of the arcopallium; LMAN, lateral magnocellular nucleus of the nidopallium; Av, avalanche; DLM, medial portion of the dorsolateral thalamic nucleus; D, dorsal; P, posterior. **(B)** Circuit diagram of the song system. Arrowheads and closed circles indicate excitatory and inhibitory connections, respectively. NC, caudal nidopallium; Arco., arcopallium; NR, rostral nidopallium; Stri., striatum. **(C)** Schematic of Serial Laser Capture RNA-sequencing (SLCR-seq). Fresh-frozen brains were cryosectioned for Laser Capture Microdissection (LCM). Individual sections of regions of interest were collected into wells of 96-well plates, and then total RNA was purified using an optimized solid phase reversible immobilization (SPRI) protocol. After RNA purification, 3’-end sequencing libraries were prepared containing unique molecular identifiers (UMI) using a custom protocol. **(D) *Left:*** Experimental overview of SLCR-seq on hearing and deaf birds. After a baseline period of song recording, birds were either deafened through bilateral cochlea removal or underwent a sham surgery. After 4, 9, or 14 days post-surgery, birds were euthanized and SLCR-seq libraries were prepared from HVC, NC, RA, Arco., LMAN, NR, Area X, and Stri. ***Right:*** Uniform Manifold Approximation and Projection (UMAP) plot of SLCR-seq data colored by section position. Each point reflects the gene expression profile of a single SLCR-seq sample. Samples show segregation by broad anatomical area — striatal (Area X), nidopallial (HVC, NC, LMAN, NR), arcopallial (RA, Arco.) — and song system nuclei from surrounding areas. **(E)** Normalized log gene expression data of three example genes — *SLC17A6*, *PVALB*, and *AR*. Each point is gene expression in a single SLCR-seq sample. *SLC17A6* is a marker for glutamatergic cells and distinctly depleted in the striatal samples; *PVALB* and *AR* are two genes known to be enriched in song system nuclei. **(F)** Coronal anatomical atlas representation of the expression of the three genes shown in panel (E). Each region is colored according to log gene expression value. D, dorsal; L, lateral.

**Supplemental Figure 2-1.**
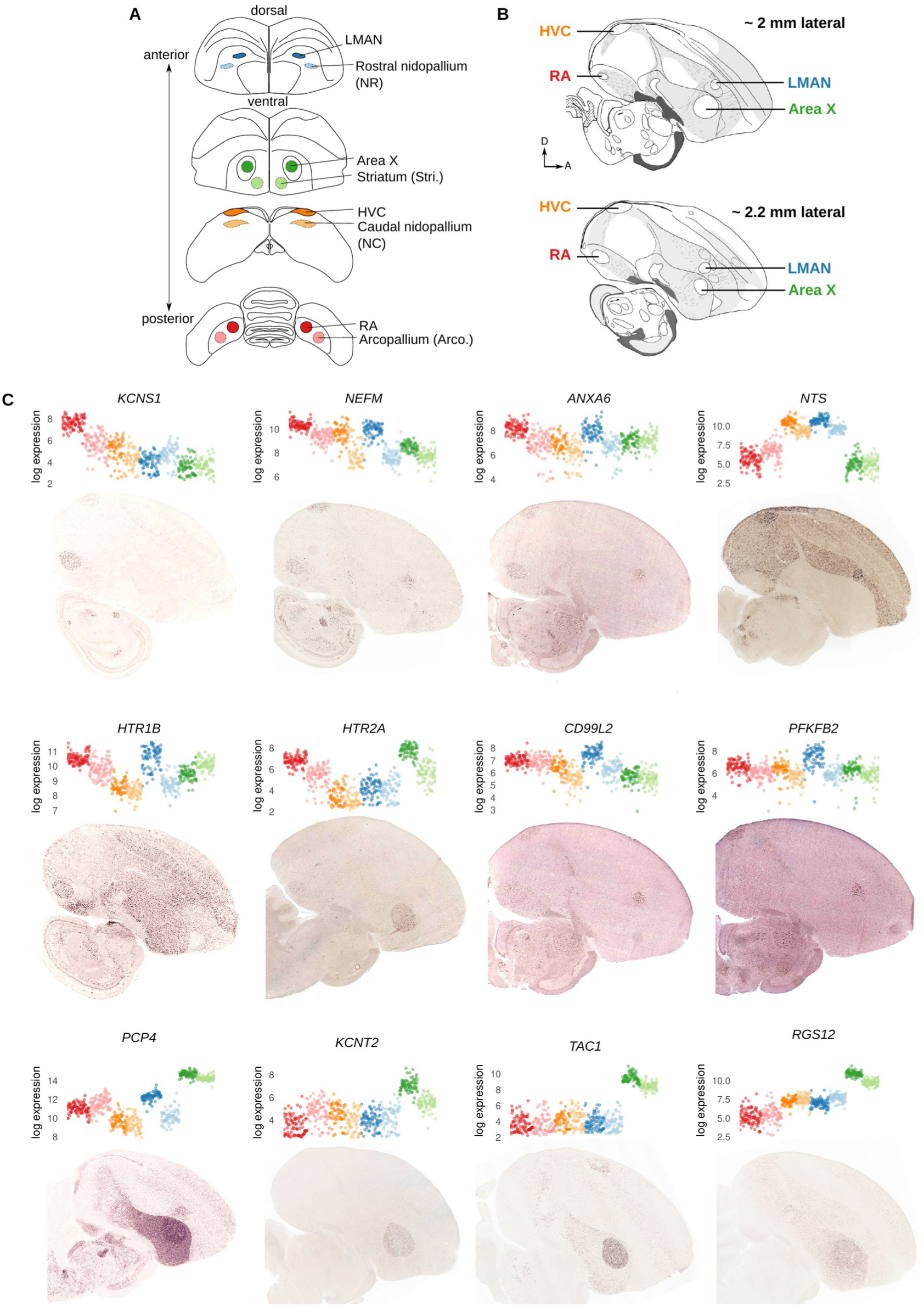
Additional validation of SLCR-seq data. Coronal sections of the finch brain showing collection locations for Serial Laser Capture RNA-seq. Each song region was identified by its position relative to anatomical landmarks, size, and characteristic Nissl staining. For each song nucleus, a region of identical size was collected outside of the nucleus in the same anatomical territory. ‘Arco.’ collection region refers to the region lateral to RA encompassing AId (dorsal part of the intermediate arcopallium) and AD (dorsal arcopallium). ‘NC’ (caudal nidopallium) collection region lies ventral to HVC outside of the HVC shelf. ‘NR’ (rostral nidopallium) collection region lies lateral to LMAN within the nidopallium. ‘Stri.’ (striatum) lies medial and ventral to Area X. **(A)** Sagittal schematics of the zebra finch brain retrieved from the ZEBrA database (http://www.zebrafinchatlas.org)(Lovell et al., 2020). Shown are two sections ~2 and ~2.2 mm lateral from the midline. Positions of RA, HVC, LMAN, and Area X are indicated. *In situ* hybridization images in panel C correspond to these representative sections. D, dorsal; A, anterior. **(C)** SLCR-seq and *in situ* hybridization expression for selected marker genes with differential expression in the song system. Each subpanel contains, at left, SLCR-seq log expression values where each point is a single SLCR-seq section. Points are colored and ordered by collection region. At right, is a representative ISH of the corresponding gene on a sagittal zebra finch brain section, obtained from the ZEBrA database.

Disruptions to song that follow the loss of auditory feedback are associated with both local alterations to song nuclei as well as to the interactions among connected components of the song system neural circuit (Brainard and Doupe, 2000; Hamaguchi et al., 2014; Kojima et al., 2013; Watanabe et al., 2006). To examine how song destabilization influences gene expression in the song system at both local and circuit levels, we developed a protocol for sample collection and RNA-seq library construction that addresses three goals: 1) precise collection of histologically defined samples, 2) ease of collecting multiple replicates per animal and brain region, and 3) reduced per-sample cost for library preparation and sequencing. This approach, termed Serial Laser Capture RNA-seq (SLCR-seq), combines the anatomical precision of laser capture microdissection (LCM) with the capacity to work with large numbers of low-input RNA samples provided by single-cell RNA-sequencing protocols (Fig. 2C, see Methods). Brains were flash-frozen without fixation then cryosectioned onto slides suitable for LCM. We visualized song nuclei using an optimized rapid Nissl staining protocol, collected single cryosections from regions of interest in 96-well plates using LCM, then purified total RNA using a custom solid phase reversible immobilization protocol. The preparation produces high quality RNA (RIN = 9.1±0.5) and yields sufficient for library preparation (one 20 μm-thick section with an area of 100,000 μm^2^ yields 1-2 ng; RA area is ~125,000 μm^2^). From this total RNA we then prepared 3’-localized RNA-seq libraries containing unique molecular identifiers adapted from protocols previously developed for single-cell RNA-sequencing (Islam et al., 2014; Kivioja et al., 2012; Picelli et al., 2014).

We used SLCR-seq to generate RNA-seq libraries for each bird from eight brain regions — HVC, RA, LMAN, and Area X, and four paired non-song regions — (Fig. 2D) with multiple LCM sections (2 to 6) collected per region per bird, yielding 598 samples after quality control filtering for the number of detected genes in each sample (mean ± s.d. number of sections per region per bird = 4.5 ± 1.5). Gene expression variation across this dataset segregated samples into three broad clusters corresponding to region of origin — arcopallium (RA and Arco.), nidopallium (HVC, NC, LMAN, and NR), and striatum (Area X and Stri.) — consistent with the different functional properties and developmental origins of these regions (Fig. 2D). Furthermore, these broad clusters were subdivided into adjacent but distinct song/non-song pairs, reflecting known similarities between song nuclei and adjacent neural regions (Nevue et al., 2020). To further validate this approach, we inspected the SLCR-seq expression values of genes with known variation across the songbird brain or enrichment in the song system (Fig. 2E,F and Figure 2-figure supplement 1). The glutamatergic neuron marker *SLC17A6* was strongly depleted in the striatal regions Area X and Stri, consistent with the relative scarcity of excitatory neurons in these regions. Genes with known variation in expression across the system, e.g. parvalbumin (*PVALB)* and androgen receptor (*AR)*, corroborated a strong correspondence between SLCR-seq expression values and *in situ* hybridization signal intensities (Lovell et al., 2020) (Fig. 2E,F and Figure 2-figure supplement 1).

### Song system-wide transcriptional signatures of song destabilization

To provide a single statistic that reflects the extent of song change for each bird, we used a previously developed method (Mets and Brainard, 2018) that builds statistical models of song in two conditions (e.g. pre and post-procedure) and calculates the distance between probability distributions generated from these models using Kullback-Leibler divergence (‘Song D_KL_’, see Methods). Here, higher values indicate larger divergence of post-procedure song compared to pre-procedure song, therefore providing a measure of song change from baseline. For each bird, we calculated Song D_KL_ between songs recorded during the two days before the procedure (deafening or sham) and those recorded on the day of euthanasia and the preceding day. Deafening resulted in a significant increase in Song D_KL_ relative to sham (Fig. 3A, hearing 0.14±0.041 log Song D_KL_ mean±sem; deaf 0.52±0.085 mean±sem, two-sided Wilcoxon rank-sum test p = 5e-4). Singing influences gene expression in the song system (Feenders et al., 2008; Horita et al., 2012; Jarvis et al., 1998; Sasaki et al., 2006; Wada et al., 2006; Warren et al., 2010; Whitney et al., 2014; Whitney and Johnson, 2005), and previous work has indicated that recent singing influences song plasticity and variability (Chen et al., 2013; Hayase et al., 2018; Hilliard et al., 2012; Miller et al., 2010; Ohgushi et al., 2015). Therefore, we also included terms for the number of songs sung on the day of euthanasia and the average number of songs sung per day in the pre-procedure period to control for constitutive differences in singing propensity. These values varied widely across birds (Fig 3B) but did not differ significantly between hearing and deafened birds (number of songs on date of euthanasia: hearing 52±18 mean±sem, deaf 55±17, two-sided Wilcoxon rank-sum test p=0.7; pre-procedure songs/day: hearing 332±39 mean±sem, deaf 339±43, two-sided Wilcoxon rank-sum test p=1).

**Figure 3.**
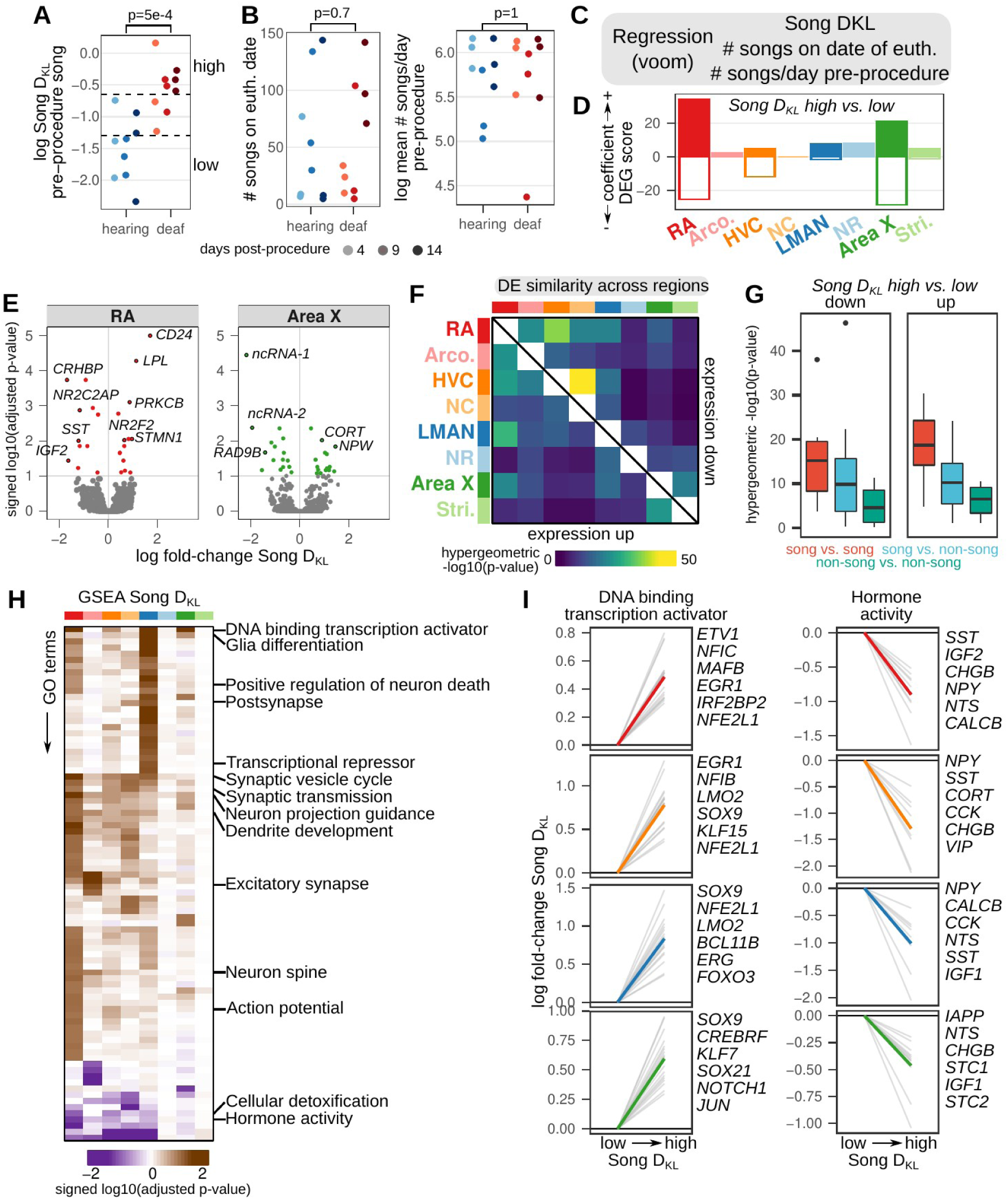
Song destabilization is associated with song system-wide alterations to gene expression. **(A)** Relative spectral distance between syllables pre- and post-procedure, represented as the mean Kullback-Leibler (KL) distance between Gaussian mixture models or ‘Song D_KL_’ (see Methods for calculation). Song DKL trends higher with increasing days from deafening. Significance calculated using a two-sided Wilcoxon rank-sum test. **(B)** Number of songs sung on euthanasia date (*left*) and log mean number of songs sung per day (*right*) for each bird grouped by hearing or deaf. Significance calculated using a two-sided Wilcoxon rank-sum test. **(C)** Differential expression analysis of song destabilization. Multiple regression using voom/limma provided estimates of gene expression fold change with variation in song deviation (Song D_KL_), number of songs on the day of euthanasia, and baseline differences in singing rate. **(D)** Differential expression gene (DEG) scores from gene expression regressions. Positive values reflect genes with increased expression, while negative values indicate genes with reduced expression. Scores are the sum of the −1 * log10(adjusted p-values) of high vs. low Song D_KL_ regression coefficients. Each score is multiplied by the sign of the coefficient to obtain a signed value. Separate coefficients were estimated for each neural region. **(E)** Volcano plots of −1 * log10(adjusted p-values) versus the log fold-change of gene expression in RA and Area X in high vs. low Song D_xKL_ birds. Signed adjusted p-values above 5 were assigned values of 5 to aid visualization. Labeled are the 10 genes with the highest signed adjusted p-value. ‘ncRNA-1’ accession is LOC116184561, ‘ncRNA-2’ accession is LOC116183441. **(F)** Similarity of song destabilization differential gene expression across the song system and surrounding regions. Heatmaps show the −log10(p-value) from hypergeometric tests comparing the expected versus observed overlap of the top 250 differentially expressed genes for each compared region, divided into genes with increased expression with song destabilization (lower left triangle) and those with decreased expression (upper right triangle). **(G)** Distribution of values from (F) comparing song versus song regions, song versus non-song regions, and non-song versus non-song regions. Box middle is the median, box upper and lower bounds are the 25th and 75th percentile, and whisker ends lie at 1.5 times the inter-quartile range. **(H)** Gene set enrichment analysis (GSEA) of song destabilization-associated genes. Shown are the Gene Ontology (GO) terms that are significant in at least one song or non-song region (adjusted p-value < 0.1, see Methods). Heatmap represents the signed log10(adjusted p-value) for each GO term and region, with the sign indicating that a given term is associated with increased or decreased expression in deaf versus hearing birds. Terms are ordered by hierarchical clustering (euclidean distance, Ward squared method). Representative terms are listed for each cluster. **(I)** Song destabilization gene expression responses of genes in two gene sets — “DNA binding transcription activator” (GO:0001216) and “Hormone activity” (GO:0005179) — that have log fold-change across song regions. Shown are the top 20 leading edge genes from GSEA and the top six of these are labeled at right.

**Supplemental Figure 3-1.**
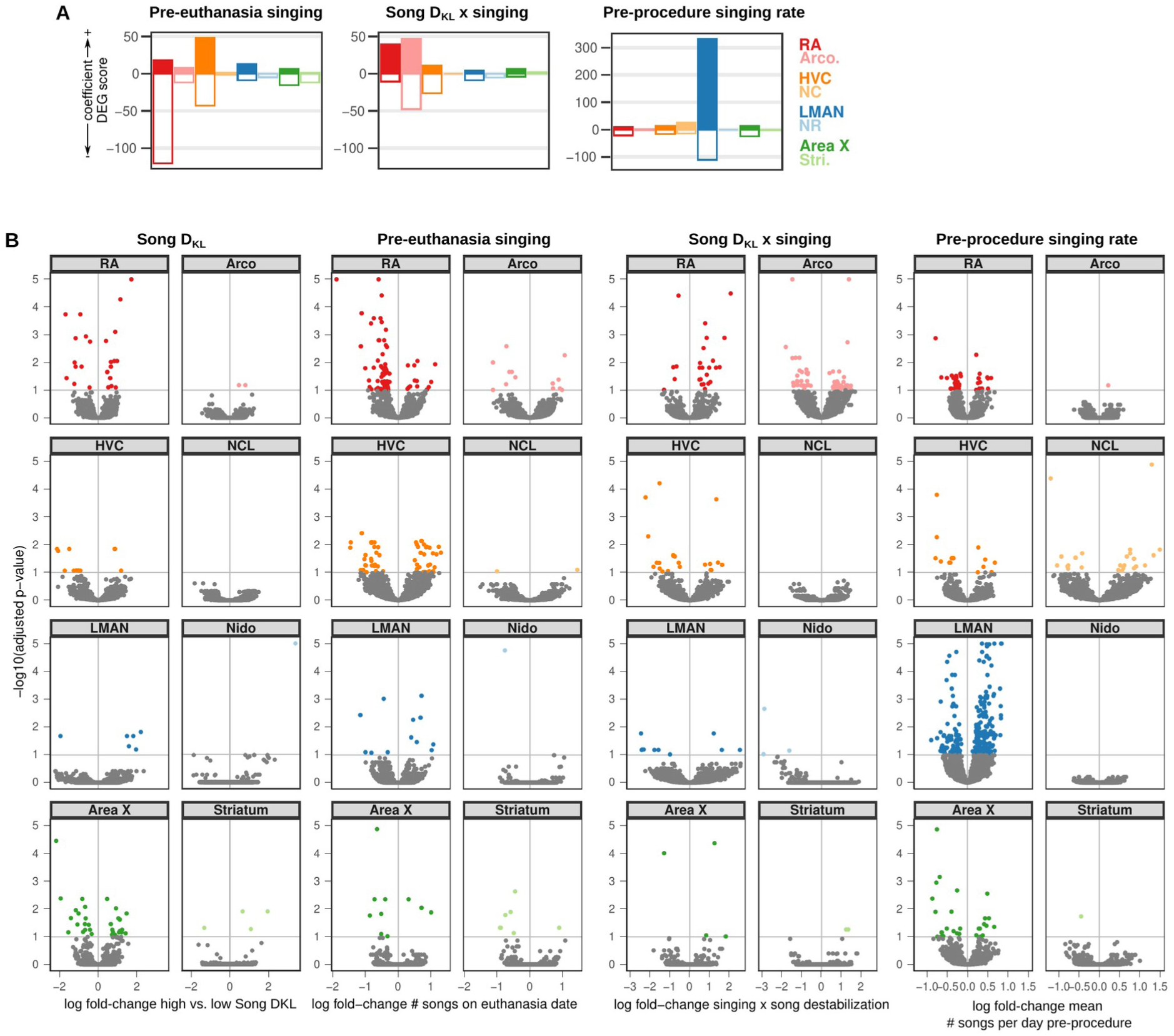
Extended analysis of song destabilization-associated gene expression. **(A)** Differential expression gene (DEG) scores from voom regression. Scores are the sum of the −1 * log10(adjusted p-values) of regression coefficients. Each score is multiplied by the sign of the coefficient to obtain a signed value. Coefficients shown are pre-euthanasia singing (# of songs sung on euthanasia date), Song D_KL_ x singing (interaction term), and pre-procedure singing rate (mean number of songs sung per day during pre-procedure days). **(B)** Volcano plots of −1 * log10(adjusted p-values) versus the log fold-change of gene expression in RA for high vs. low Song D_KL_ (‘Song D_KL_’), the number of songs sung on the day of euthanasia (‘Pre-euthanasia singing’), the interaction between Song D_KL_ and the number of songs sung on the day of euthanasia (‘Song D_KL_ x singing’), and the mean number of songs sung per day during pre-procedure days (‘Pre-procedure singing rate’). Signed adjusted p-values above 5 were assigned values of 5 to aid visualization.

We used multiple regression to identify genes whose expression varied with song destabilization (Song D_KL_) (Fig. 3C). A subset of hearing and deaf birds showed overlapping Song D_KL_ values (3 birds in each condition). To detect genes with expression differences associated with song destabilization, we compared birds in each group that had Song D_KL_ values outside of this overlapping range (Fig 3A, 6 hearing birds with ‘low’ song DKL values, and 6 deaf birds with ‘high’ song DKL values). To identify regional patterns of differential expression, we used a differential expression gene (DEG) score that incorporates the number of differentially expressed genes and their adjusted p-values (Fig. 3D and Supplemental Table 2, see Methods). Song regions generally showed greater expression differences than non-song regions for both Song D_KL_ and singing-rate (Fig. 3D and Figure 3-figure supplement 1). Across the song regions, the largest changes to gene expression between high and low Song D_KL_ occurred for the forebrain motor output nucleus RA and the striatal component of the song system Area X (Fig. 3D, red and green bars). For these regions, the most significantly modulated genes (adjusted p-value < 0.1) were equally likely to be upregulated versus downregulated in deafened versus hearing birds (Fig. 3E; for RA, 14 genes upregulated and 11 genes downregulated; for Area X, 15 genes up, 16 genes down). Among the most highly upregulated genes in RA were the plasticity-associated gene protein kinase C β (*PRKCB*) (Chu et al., 2014; Fioravante et al., 2014), whose protein levels were previously shown to be upregulated in RA following deafening (Watanabe et al., 2002); the microtubule-destabilizing protein stathmin 1 (*STMN1*), which has roles in long term potentiation and fear memory formation (Shumyatsky et al., 2005); *CD24*, a surface protein that influences neurite extension (Gilliam et al., 2017); and the lipid processing enzyme lipoprotein lipase (*LPL*), which is implicated in memory formation and Alzheimer’s disease pathology (Wang and Eckel, 2012; Yu et al., 2015). Likewise, among the most downregulated genes in RA were secreted neuromodulatory proteins including corticotropin-releasing hormone binding protein (*CRHBP*), somatostatin (*SST*), and insulin growth factor 2 (*IGF2*), which each have described roles in regulating neuronal physiology and neural plasticity (Chen et al., 2011; Li et al., 2016; Song et al., 2021).

To further examine the neural-circuit wide structure of gene expression across the song system and surrounding regions, we pairwise intersected the lists of the top Song D_KL_ differentially expressed genes (250 genes with the lowest p-values) for each region and calculated the degree of overlap using a hypergeometric test (Fig. 3F and G). Song destabilization-associated differential gene expression was more similar between song regions than between both song and non-song pairs and non-song regions with each other (Fig. 3G), indicating that the song system exhibits, in part, a common transcriptional response to song destabilization that is not shared in adjacent regions. We performed gene set enrichment analysis (GSEA) of differentially expressed genes to identify pathways that show coherent gene expression responses to song destabilization (Fig. 3H and Supplemental Table 3). Several pathways exhibited similar expression responses across all four song regions, including those related to transcription regulation, glia differentiation, and hormone activity (Fig. 3H and I). Genes related to synaptic transmission were differentially expressed across multiple pallial regions, including song regions RA, HVC, and LMAN as well as the non-song region NCL. Neuron spine-associated genes were upregulated across RA, Arco., and HVC, consistent with previous reports of altered spine dynamics in the song motor pathway following deafening (Peng et al., 2013, 2012a; Tschida and Mooney, 2012; Zhou et al., 2017).

### Correlated gene modules associated with song destabilization

The regression analysis above identified differential expression at the level of individual genes but may have missed subtler expression responses that are correlated across multiple genes. To better identify groups of differentially expressed genes with similar responses to song destabilization, we leveraged the large sample numbers of the SLCR-seq dataset to perform gene-gene correlation network analysis for each region separately using MEGENA (Multiscale Embedded Gene Co-expression Network Analysis), an approach that generates sparse networks of covarying genes by applying a topological constraint to co-expression networks (Song and Zhang, 2015) (Fig. 4A and Figure 4-figure supplement 1). Using this method, we constructed gene-gene correlation networks for each song and non-song region separately, combining SCLR-seq samples across all birds, both hearing and deafened. Mapping song destabilization fold-change of each gene onto the RA network showed a segregation between genes with increased and decreased expression, indicating that expression differences associated with song destabilization state are prominent drivers of network structure (Fig. 4B). This segregation was also seen for the HVC, LMAN, and Area X networks (Figure 4-figure supplement 1A). MEGENA employs a hierarchical module detection algorithm to identify correlated sets of genes at different levels of resolution (Supplemental Table 4). For each module in each region’s network, we averaged high-vs-low Song D_KL_ regression coefficients for its member genes and compared these observed mean values to a shuffled distribution of mean values, generated by sampling the same number of module genes across the network at random. Several modules in each region’s network showed significantly higher or lower mean fold-change relative to a distribution of mean fold-changes from sets of randomly selected genes (100 random samplings of genes from the network, shuffled p-value < 0.01, Fig. 4C and Figure 4-figure supplement 1B).

**Figure 4.**
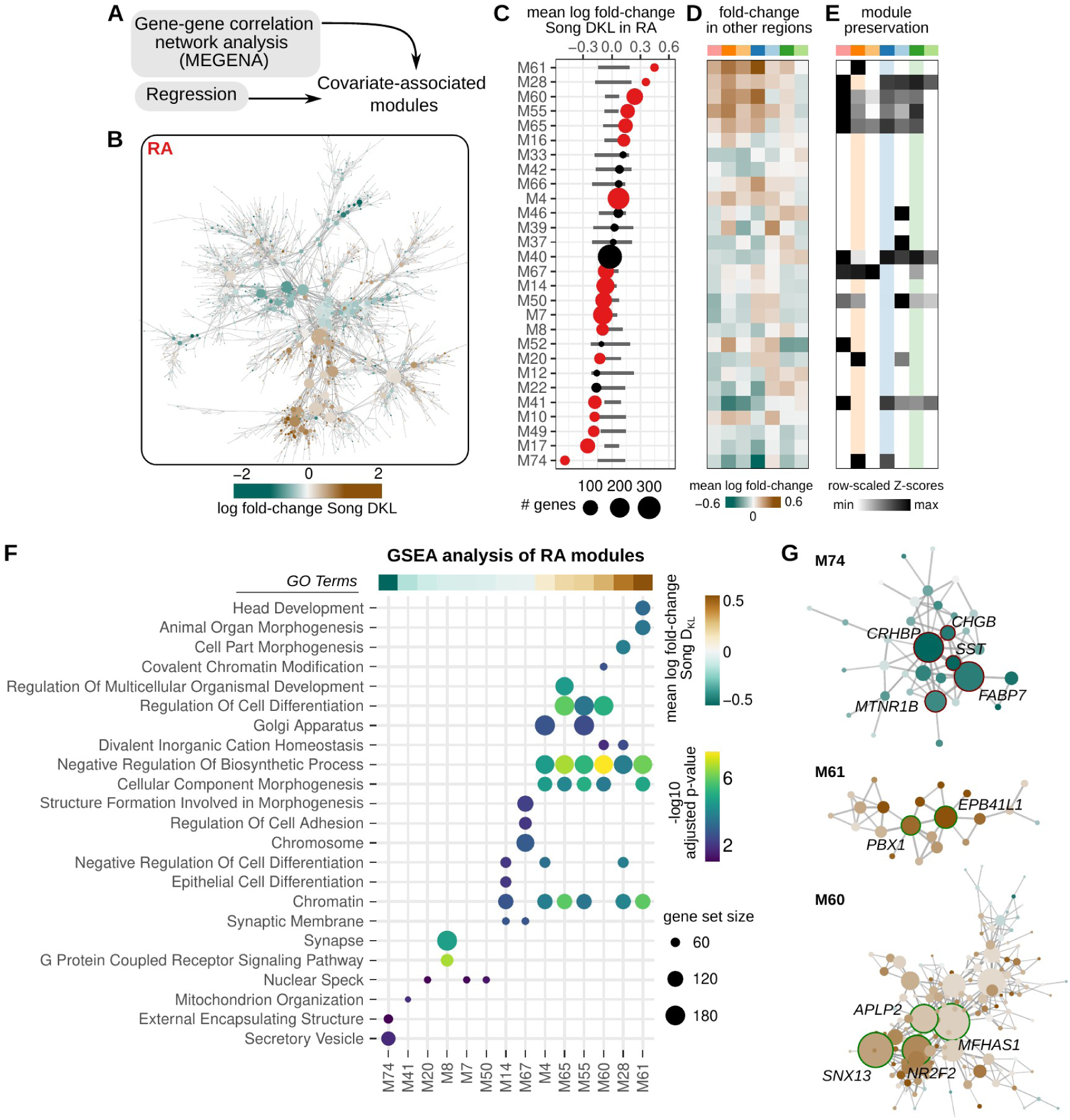
Correlated modules of gene expression associated with song destabilization. **(A)** To identify correlated patterns of gene expression, gene-gene correlation networks were constructed for each region using MEGENA (Multiscale Embedded Gene Co-expression Network Analysis). These networks were then used to identify correlated sets of gene modules. Estimated regression coefficients were mapped onto correlation networks to identify covariate associated expression modules. **(B)** Gene-gene correlation network for RA. Each node is colored by the log fold-change expression between deaf and hearing birds. **(C)** Average song destabilization gene expression change for each RA module. Error bars are null distributions generated by repeatedly sampling the network (100 times) for the number of nodes in a given module then averaging their high vs. low Song D_KL_ fold-changes. Dots that are colored have mean coefficient values that are lower or higher than 1% or 99% of the sampled distribution, respectively. **(D)** Average change in expression of RA modules across each song system and non-song system region. **(E)** Preservation scores for RA modules in the correlation networks of other song and non-song system regions. Only significant values are shown (Bonferroni p-values < 0.01), and values are scaled to the maximum and minimum for each module to show relative levels of preservation across regions. **(F)** Gene set enrichment analysis (GSEA) of RA modules with significant gene expression alteration with song destabilization. Mean fold-change values for each module, as represented in Fig. 4B, are shown at the top of the GSEA plot. Shown are at most the top 5 significant Gene Ontology (GO) terms (GSEA adjusted p-values < 0.2). **(G)** Network diagrams for three modules, M61, M60, and M74, that show large deviations with song destabilization. Labeled and highlighted are selected hub genes for each module (see Methods for classification). Node colors indicate log fold-change expression between deaf and hearing birds (scale given in Figure 4A).

**Supplemental Figure 4-1.**
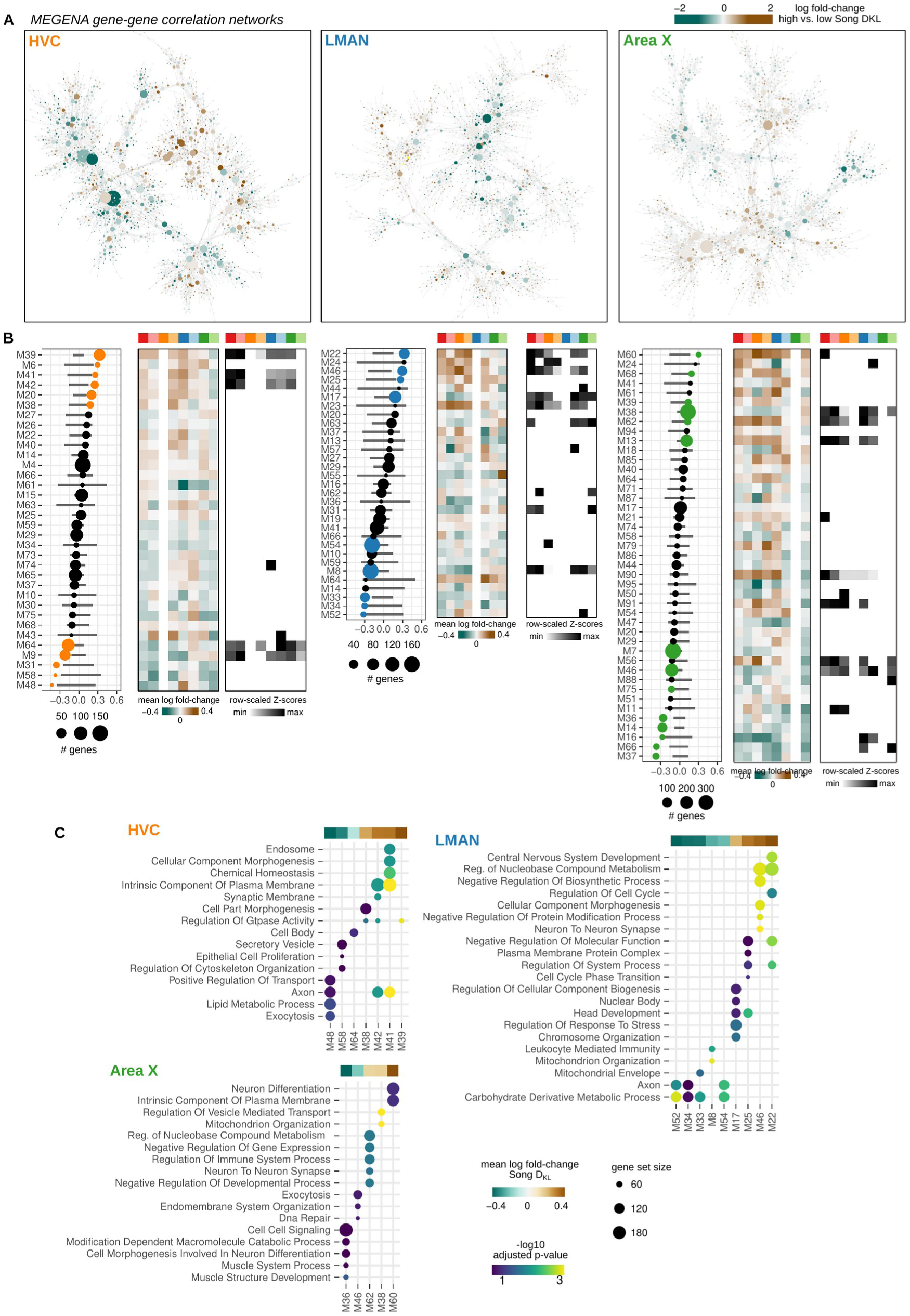
Network analysis of song destabilization-associated gene expression. **(A)** MEGENA gene-gene correlation network for HVC, LMAN, and Area X. Each node is colored by the log fold-change expression between high and low Song D_KL_ groups. **(B)** Average differential expression (**left**), patterns of differential expression across regions (**middle**), and preservation scores (**right**) for the modules identified in the HVC, LMAN, and Area X networks. Analysis and figure presentation is as described in Figure 4C-E. **(C)** Gene set enrichment analysis (GSEA) of HVC, LMAN, and Area X modules with significant gene expression alteration with song destabilization. Mean fold-change values for each module, as represented in Fig. S4B, are shown at the top of the GSEA plot. Shown are at most the top 5 significant Gene Ontology (GO) terms (GSEA adjusted p-values < 0.2).

To assess the similarity between modules in the correlation networks of one brain region and those in the networks of other brain regions, we calculated a module preservation score (see Methods) and found that RA destabilization-associated modules were preserved to different degrees in networks for the other song regions. In addition, several RA modules showed similar response patterns in other song regions, such that modules upregulated in RA were upregulated in HVC, LMAN, and Area X and likewise for downregulated modules (Fig. 4D,E and Figure 4-figure supplement 1B). This pattern is consistent with the overall similarity in differential expression seen among song regions using the regression analysis above (Fig. 3F). Gene set enrichment analysis indicated that differential RA modules are enriched for a range of biological pathways (Fig. 4F and Figure 4-figure supplement 1C). Notably, the top downregulated module (M74) was enriched for secreted proteins, such as *CRHBP*, *SST*, and *CHGB* (Fig. 4G). Upregulated modules were enriched for genes involved in development, morphogenesis, and gene regulation, including *PBX1, NR2F2, ZHX3*, and *ANKRD11*.

### Cell type expression of song destabilization-associated genes

After establishing a circuit-wide view of gene expression responses to song destabilization, we investigated the cellular specificity of these responses to understand what cell classes exhibit the most substantial transcriptional changes and may play a role in deafening-induced song plasticity. To do so, we integrated the above SLCR-seq data with a previously generated single-nucleus and single-cell RNA-sequencing dataset from HVC and RA of hearing adult male finches (Colquitt et al., 2021) (Fig. 5A). In that work, we compared songbird neuronal classes in HVC and RA to those in mammals and identified a high degree of transcriptional similarity across several neuronal classes (Fig. 5B).

**Figure 5.**
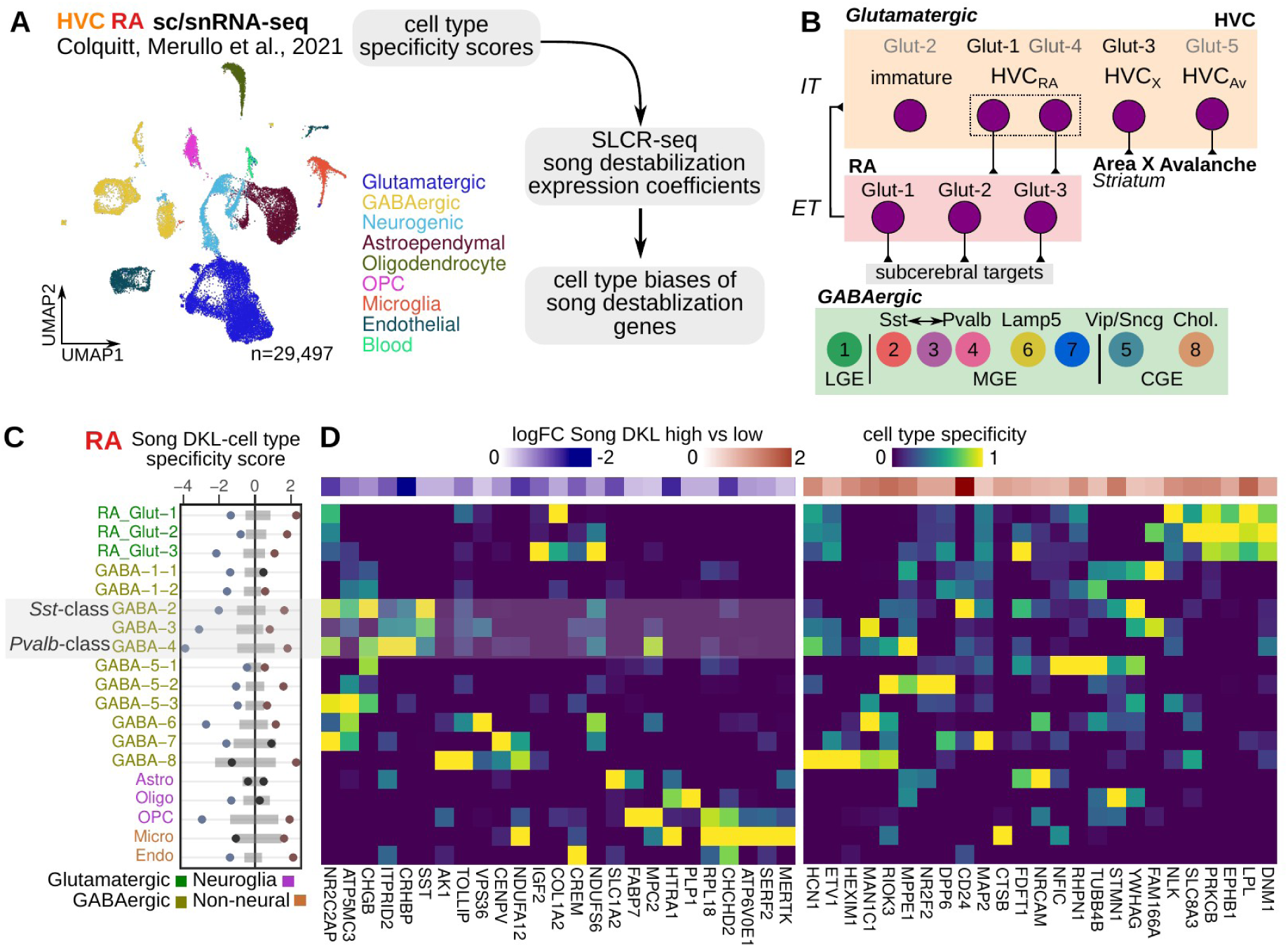
Cell-type specificity of destabilization modulated genes. **(A)** Schematic of the approach to determine the cell type expression biases of genes that are differentially regulated with song destabilization. A previously generated cell-resolved gene expression dataset for RA and HVC (Colquitt et al., 2021) was combined with RA and HVC song destabilization regression coefficients from this study to compute a cell type bias score (see Methods). Shown also is a UMAP plot of the full dataset with major cell type groups indicated. OPC, oligodendrocyte precursor cell. **(B)** Schematic of neuronal cell types in the song motor pathway, as previously defined in (Colquitt et al., 2021). HVC glutamatergic neurons are broadly similar to intratelencephalic (IT) mammalian neocortical neurons from multiple layers, and RA neurons are similar to extratelencephalic (ET) neurons from layer 5. Eight primary GABAergic clusters are found equally in both HVC and RA and are organized into clusters corresponding to subpallial regions of origin — lateral, medial, and caudal ganglionic eminences (LGE/MGE/CGE). The LGE-class GABA-1 has no known correspondence with mammalian neocortical neurons; GABA-2 is transcriptionally similar to Sst-class neurons, GABA-4 is similar to Pvalb-class neurons, and GABA-3 is transcriptionally intermediate between GABA-2/4. **(C)** Integration of cell type specificity scores and song destabilization differential expression to identify cell type-associated transcriptional effects of song destabilization. For the top 50 differentially expressed genes, cell type specificity was multiplied by log fold-change between high and low Song DKL birds. These values were then split by sign then summed within each cell type to yield a cell type Song DKL score. Gray bars indicate the distribution (1% to 99%) of Song D_KL_-cell type specificity scores for 100 random sets of 50 genes. **(D)** RA cell type specificity scores for top Song D_KL_ differentially expressed genes, divided into upregulated and downregulated genes. Values are scaled for each gene such that the cell type with the highest specificity score equals 1 and that with the lowest equals 0. At the top of each specificity score heatmap is the log fold-change expression for high vs. low Song D_KL_.

**Supplemental Figure 5-1.**
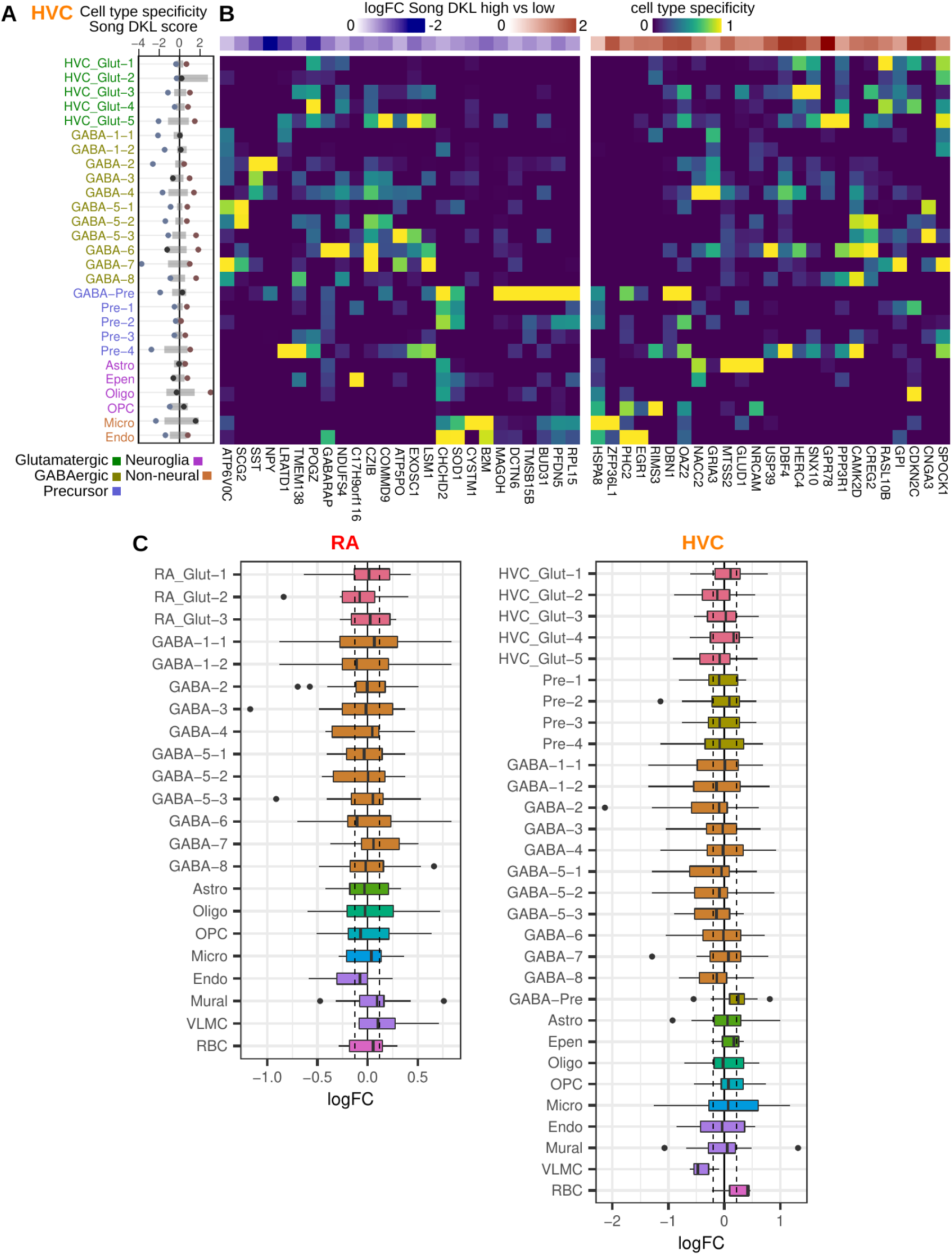
Cell type-associated differential expression. **(A)** Integration of cell type specificity scores and Song D_KL_ coefficients to identify cell type-associated transcriptional effects of song destabilization in HVC. For the top 50 differentially expressed genes, cell type specificity was multiplied by its log fold-change between high and low Song D_KL_ birds. These values were then split by sign then summed within each cell type to yield a cell type Song D_KL_ score. Gray bars indicate the distribution (1% to 99%) of Song D_KL_-cell type specificity scores for 100 random sets of 50 genes. **(B)** HVC cell type specificity scores for top Song D_KL_ differentially expressed genes, divided into upregulated and downregulated genes. Values are scaled for each gene such that the cell type with the highest specificity score equals 1 and that with the lowest equals 0. At the top of each specificity score heatmap is the log fold-change expression for high vs. low Song D_KL_. **(C)** Distributions of Song DKL high vs. low log fold-change for the top 25 marker genes for each cell type. Dashed lines indicate the 1% and 99% median log fold-change for 100 randomly selected sets of 25 genes.

For each gene, we computed a cell type destabilization score — the product of a gene’s cell type specificity with its fold-change between high and low Song D_KL_ — to assay cellular biases of destabilization-associated expression (Fig. 5C and D, and Figure 5-figure supplement 1A and B). In RA, which showed the strongest transcriptional changes as described above (Fig. 3D), differentially expressed genes were most strongly localized to neurons. In particular, genes with reduced expression during song destabilization, such as *CRHBP*, *SST*, *NPY*, and *CHGB,* showed a bias toward *Sst*- and *Pvalb*-class interneurons (GABA-2/3/4). In addition, several upregulated genes, such as *PRKCB, EPHB1,* and *DNM1*, were biased toward RA glutamatergic neurons. HVC showed a similar pattern of cell-type expression, with genes that had reduced expression biased toward *Sst*-class interneurons as well as LGE-class GABA-1 and MGE-class GABA-7 interneurons (Figure 5-figure supplement 1A,B). These cellular expression biases could arise from increases or decreases in the abundances of defined cell populations. To determine if these cellular biases do reflect a bulk loss of particular cell types, we analyzed the differential expression of marker genes for each cell type (Figure 5-figure supplement 1C). In both song nuclei, markers for each neuronal cell type (see Methods for definition) showed no significant difference between high and low Song D_KL_ conditions (median of fold-changes greater than 99% or less than 1% of medians from randomly selected genes), indicating that the cell type biases of destabilization associated-genes are not due to changes in cell type abundance, but rather to the expression levels of specific genes within defined cell classes.

### Inter-region correlation of gene expression is reduced in deafened birds

The foregoing analyses focused on comparing gene expression responses to deafening that are local to each region. However, the song system is an interconnected neural circuit and gene expression in one region could be correlated with that in others due to shared patterns of neural activity, common responses to hormonal signaling, or to baseline expression differences across regions that vary in a concerted fashion across individuals. By similar logic, manipulations such as deafening that could disrupt global patterns of neural or hormonal signaling might result in alterations in the patterns of inter-region correlations in gene-expression.

To determine whether and how deafening alters inter-region correlations in gene expression, we first identified genes that have correlated expression between brain regions across birds. Briefly, for each gene, we calculated correlation values for each pairwise combination of brain regions, yielding region-by-region correlation matrices (Fig. 6A). We identified significant correlations as those that were less than the 2.5% quantile or greater than the 97.5% quantile of a shuffled distribution (see Methods). We calculated the across-bird gene expression similarity between regions as the number of thresholded correlations. This analysis revealed several notable relationships among brain regions. First, each song nucleus had the highest gene correlation strength with its paired non-song region (with the exception of HVC which had generally weaker correlation strengths with other regions), consistent with the shared molecular profiles of each song nucleus with surrounding tissue. Second, the nuclei of the vocal motor pathway (HVC and RA) and anterior forebrain pathway (LMAN and Area X) were more correlated with each other than with nuclei in the other pathway. Third, normalizing correlation strength for each song nucleus recovered known connections between nuclei (Fig. 6B): HVC displayed strong correlations with its target RA, and LMAN was strongly correlated with both of its direct targets, RA and Area X. Interestingly, we found relatively weaker gene correlation strength between HVC and its target Area X.

**Figure 6.**
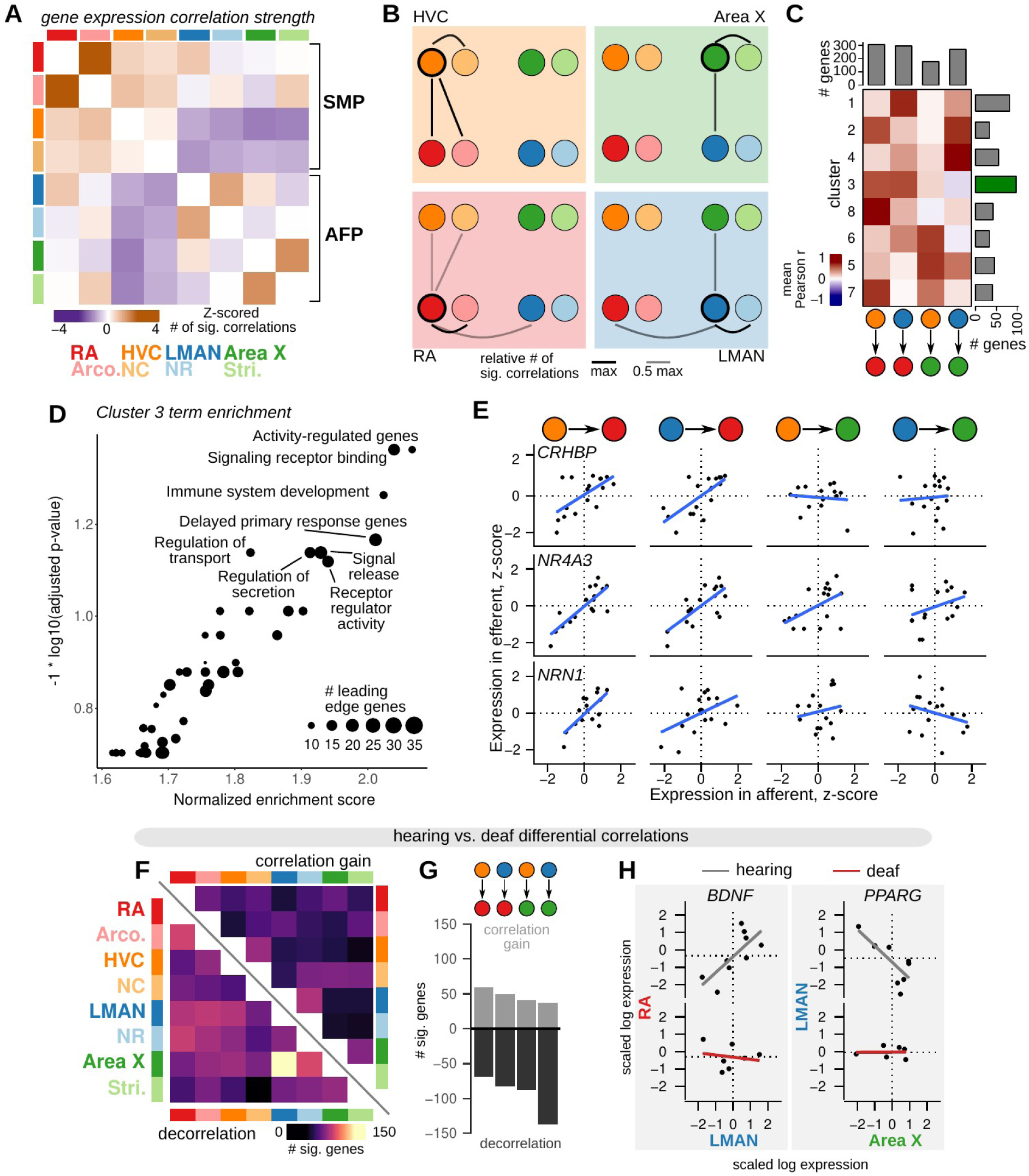
Inter-region gene expression correlation and decorrelation with song destabilization. **(A)** Inter-region gene expression correlations across song and non-song regions. For each region, the 500 genes with the highest variability across birds were selected (see Methods), then the expression of each gene was correlated across regions. Significant genes were called as those with an observed correlation less than 1% or greater than 99% of a shuffled correlation distribution (100 shuffles, calculated for each pairwise comparison between regions). **(B)** Representation of the data in (A) showing the strength of gene expression correlation between each song system nucleus and other assayed regions. **(C)** Patterns of inter-region gene expression correlations. Genes were clustered into 8 clusters by their pairwise correlation values between HVC-RA, LMAN-RA, HVC-Area X, and LMAN-Area X. Heatmap shows mean correlations within each cluster (rows) and region comparison (columns). Barplots represent the number of genes in each row or column. Highlighted is cluster 3, the HVC-RA and LMAN-RA correlation cluster, which has the greatest number of genes. **(D)** GSEA indicates that cluster 3 is enriched for genes that are activity-dependent and have signaling-related function. **(E)** Expression of three cluster 3 genes across pairs of song system regions with direct projections — HVC to RA, LMAN to RA, HVC to Area X, and LMAN to Area X. Each point is the z-scored expression estimate for each nucleus in one bird. **(F)** Analysis of inter-region gene expression correlations compared between hearing and deaf birds. For each gene, region pair, and condition, a Pearson correlation was calculated, then differential correlation was calculated as the difference between unsigned correlations for hearing versus deaf conditions. To determine significance, 100 random permutations of the expression data were made for each gene, region pair, and condition and differential correlation was computed as above. Genes with observed differential correlations in the top or bottom 2.5% of the shuffled distribution were considered significant. Heatmap shows the number of genes that became decorrelated (bottom-left) or gained correlation (top-right) in deaf versus hearing birds. **(G)** Total number of genes that show significant correlation gain or decorrelation in the four pairs of assayed song system regions with direct projections. **(H)** Examples of genes that show decorrelation in deafened birds relative to hearing birds. *BDNF* (brain derived neurotrophic factor) expression is correlated between RA (y-axis) and LMAN (x-axis) in hearing birds but shows no inter-region correlation in deafened birds. Similarly, *PPARG* expression is correlated between LMAN (y-axis) and Area X (x-axis) in hearing birds but uncorrelated in deaf birds. Each point is the z-scored expression estimate for one bird.

To identify genes that have shared patterns of inter-region correlation across multiple song nuclei, we next clustered genes by the similarity of the correlations between song nuclei known to be directly connected, HVC-RA, LMAN-RA, HVC-X, and LMAN-X (Fig. 6C). This analysis generated a diversity of patterns with most genes showing correlated expression among the three pallial song nuclei, HVC-RA and LMAN-RA (cluster 3). Gene set enrichment analysis indicated that this cluster is enriched for genes that are associated with signaling receptor binding and that are responsive to neural activity (Fig. 6D). Indeed, the genes most strongly associated with HVC-RA and LMAN-RA correlations included the activity-dependent genes *CRHBP*, *NR4A3,* and *NRN1* (Fig. 6E).

We then assessed how deafening alters gene expression correlations across the song system. To do so, we computed pairwise correlations for each gene between each region for hearing and deaf birds separately, then computed a differential matrix comparing absolute correlations in deaf birds to those in hearing birds (Fig. 6F-H). Differentially correlated genes were defined as those with a deaf versus hearing value less than (decorrelation) or greater than (correlation gain) the extreme values of a shuffled distribution calculated for each pairwise comparison (2.5% or 97.5%, respectively). Overall, each directly connected pair of song regions had a greater number of genes with reduced correlation in deaf versus hearing birds than increased correlation (Fig. 6 F, G and Supplemental Table 5). Two of the most strongly decorrelated genes highlight this effect. Expression of the neurotrophic factor *BDNF* was positively correlated between LMAN and RA in hearing birds but was uncorrelated in deaf birds; similarly, expression of the nuclear receptor *PPARG* was negatively correlated between LMAN and Area X in hearing birds but was uncorrelated in deaf birds (Fig. 6H).

### Loss of afferent input to the motor pathway affects expression of song destabilization-associated genes

The output nucleus of the anterior forebrain pathway, LMAN, is required for adaptive plasticity to song and moment-by-moment song variability and is one of the two major afferents to the motor nucleus RA (Andalman and Fee, 2009; Kao et al., 2005; Nottebohm et al., 1982; Olveczky et al., 2005; Warren et al., 2011; Williams and Mehta, 1999) (Fig. 2A,B). Lesions of this nucleus result in reduced song variability (Kao and Brainard, 2006) and, when performed before cochlea removal, prevent song destabilization (Brainard and Doupe, 2000), indicating that deafening generates plasticity signals that require inputs from LMAN. We hypothesized that lesions of LMAN would establish a molecular state in RA similar to that found in other low variability and low plasticity conditions, such as that in normal hearing adult birds (versus deaf birds). To assess LMAN’s influence on gene expression in the song motor pathway, we unilaterally lesioned LMAN in five adult male birds (Fig. 7A and Figure 7-figure supplement 1A-C). Unilateral LMAN lesions did not grossly alter song, and song stability as measured by Song D_KL_ was similar to that for unlesioned hearing birds (Figure 7-figure supplement 1D). Sixteen days after lesioning, we collected HVC, NCL, RA, Arco., and the primary auditory area Field L for SLCR-seq (Fig. 7B, 91 libraries total). Song regions from each hemisphere were collected independently to allow within bird comparisons between regions ipsi- and contralateral to the lesion. LMAN was not substantially lesioned in one bird (lesion extent ~0% of LMAN volume, see Methods), and samples from each hemisphere for this bird were treated as unlesioned. Unlike mammals, birds do not have an interhemispheric connection at the level of the forebrain, such that there is no direct connectivity between song system nuclei across hemispheres (Nottebohm et al., 1982, 1976). We reasoned that gene expression modulated directly by LMAN activity would show specific effects in its direct target RA relative to regions that do not receive direct afferents from LMAN such as HVC and surrounding regions that are not part of the song system (Arco., NCL, and Field L).

**Figure 7.**
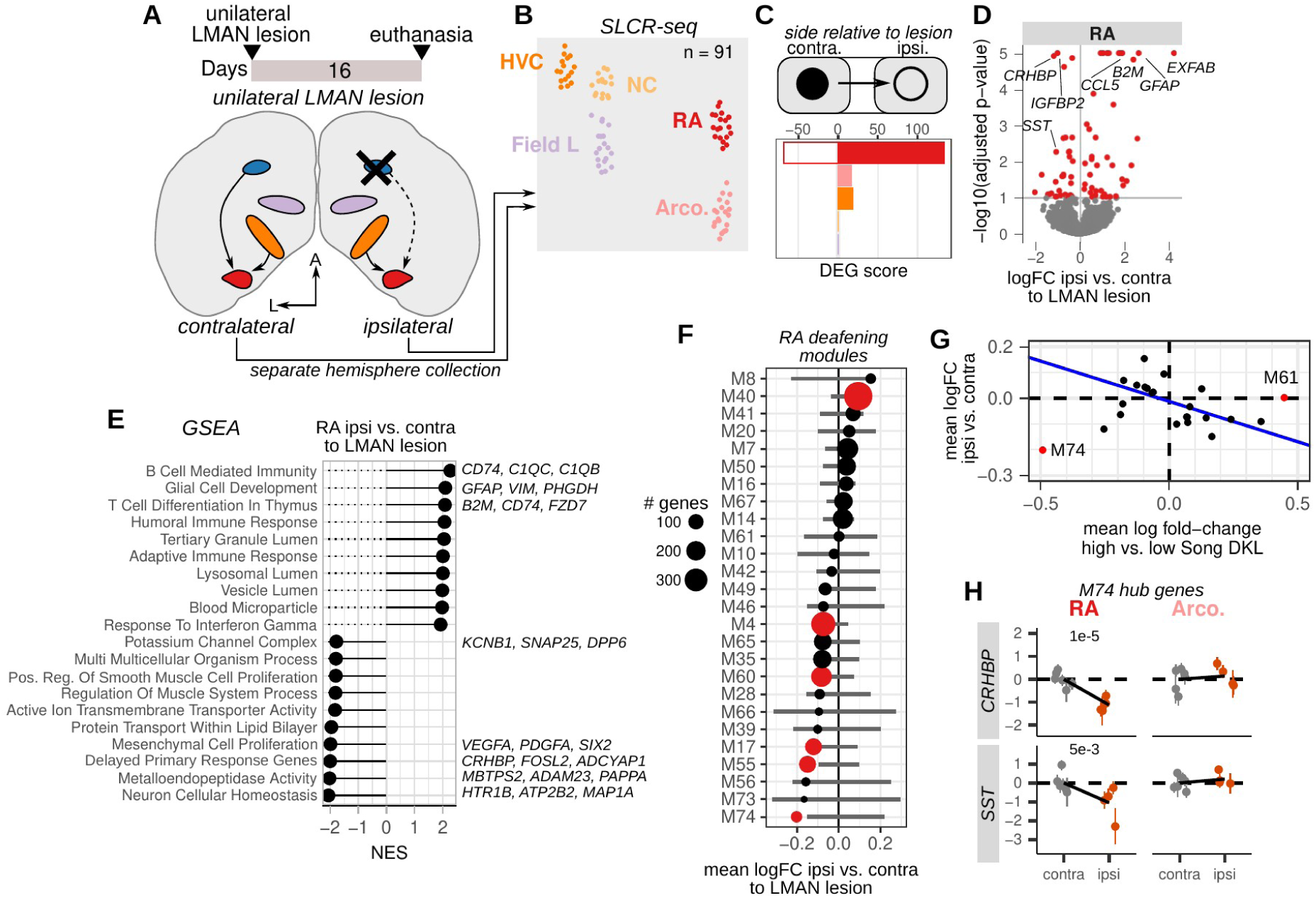
Loss of afferent input to the motor pathway nucleus RA alters destabilization-associated gene expression. **(A)** Schematic of unilateral LMAN lesions and sample collection for SLCR-seq. Five birds received unilateral LMAN lesions (3 left and 2 right hemisphere). After 16 days, birds were euthanized, and HVC, NC, RA, Arco., and the primary auditory region Field L were collected for SLCR-seq. Each hemisphere was processed separately to examine the ipsilateral versus contralateral influence of LMAN lesioning on gene expression in hearing birds. **(B)** UMAP plot of SLCR-seq samples colored by region. **(C)** Within-bird differential expression analysis of the influence of LMAN lesions on different regions of the songbird brain. Shown are differential expression gene (DEG) scores for each region assayed. Positive values reflect genes with increased expression, while negative values indicate genes with reduced expression. DEG scores are calculated as the sum of the −1 * log10(adjusted p-values) of regression coefficients for gene expression in brain regions ipsilateral versus contralateral to the LMAN lesion. Each score is multiplied by the sign of the coefficient to obtain a signed value. Separate coefficients were estimated for each neural region. **(D)** Volcano plot showing the genes that had the most significant difference in expression (red points) between RA on the ipsilateral versus contralateral side of LMAN lesion, quantified as −1 * log10(adjusted p-values) versus the log fold-change of gene expression for RA ipsilateral versus contralateral to the LMAN lesion. Signed adjusted p-values above 5 were assigned values of 5 to aid visualization. **(E)** Gene set enrichment analysis of differential expression in RA ipsilateral versus contralateral to the LMAN lesion side. Top leading edge genes are listed at right. NES, normalized enrichment score. **(F)** Average expression change between ipsilateral and contralateral RA for each Song DKL module identified in Figure 3. Error bars are null distributions generated by repeatedly sampling the network (100 times) for the number of nodes in a given module then averaging their differential ipsilateral versus contralateral coefficients. Dots are colored that have mean coefficient values that are lower or higher than 1% or 99% of the sampled distribution, respectively. **(G)** Comparison of mean fold-change expression differences in RA between high-vs-low Song D_KL_ and contralateral versus ipsilateral to LMAN lesions. Blue line indicates linear regression through the data after excluding outlier modules M74 and M61. **(H)** Expression of two M74 hub genes, *CRHBP* and *SST*, between RA and Arco contralateral or ipsilateral to the LMAN lesion. Each dot is the estimated gene expression within a given bird and region, and error bars are standard errors of this estimate. Adjusted p-values were obtained from the ipsilateral versus contralateral regression analysis.

**Supplemental Figure 7-1.**
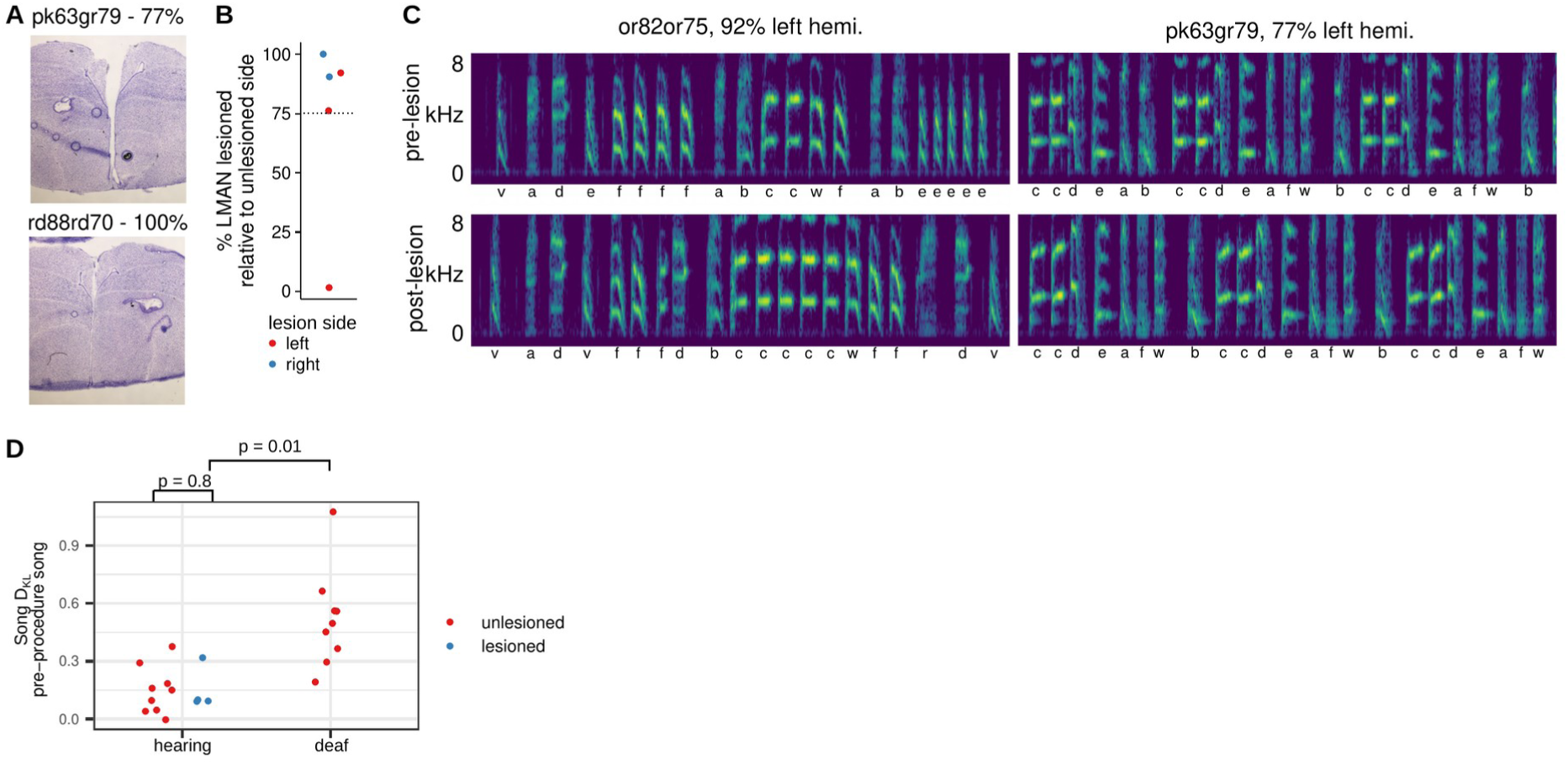
Validation and quantification of unilateral LMAN lesions. **(A)** Nissl stains of coronal sections from two example birds with unilateral LMAN lesions in the left hemisphere (bird identification pk63gr79) and right hemisphere (rd88rd70). Percentages indicate the fraction of LMAN lesioned, relative to the size of LMAN on the unlesioned side (see Methods for calculation). **(B)** LMAN lesion percentages across birds colored by lesion side. Dashed horizontal line indicates the threshold used to consider LMAN as lesioned. **(C)** Example spectrograms from two birds before and after unilateral LMAN lesion. Percentages indicate LMAN lesion extent as indicated in panels A and B. Lettering below the spectrograms are manually assigned syllable labels. **(D)** Song D_KL_ of unlesioned hearing and deaf birds (from main deafening dataset) and hearing birds with unilateral LMAN lesion. P-values are from two-sided Wilcoxon rank-sum tests.

**Supplemental Figure 7-2.**
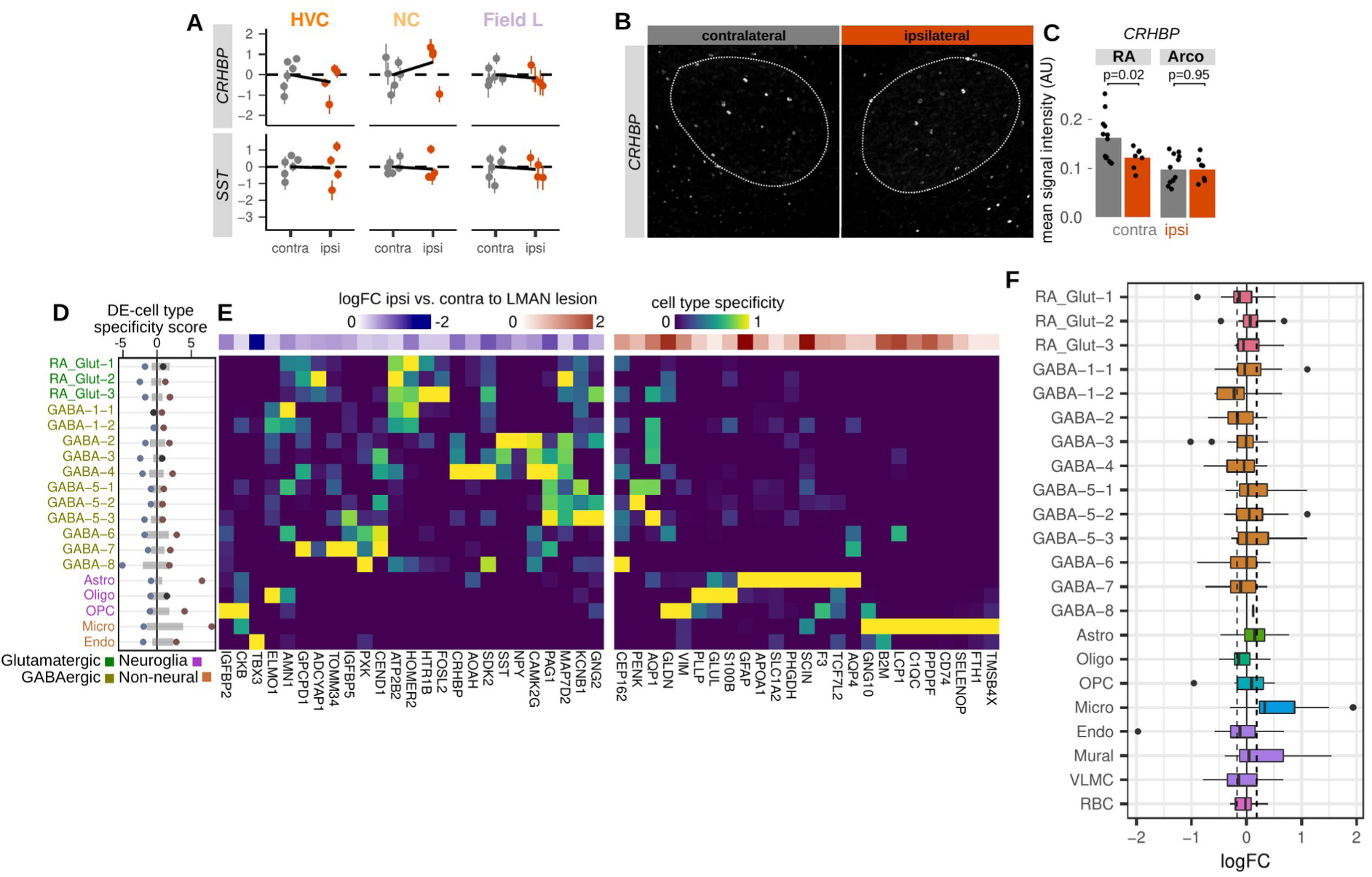
Validation of unilateral LMAN lesion effects on RA gene expression. **(A)** Expression of two M74 hub genes, *CRHBP* and *SST*, between HVC, NC, and Field L contralateral or ipsilateral to the LMAN lesion. Each dot is the estimated gene expression within a given bird and region, and error bars are standard errors of this estimate. **(B)** *In situ* hybridizations showing expression of *CRHBP* in RA either ipsilateral or unilateral to LMAN lesion. **(C)** Quantification of *CRHBP* signal intensities. Each point is the average ISH intensity in each RA or surrounding arcopallium (lesioned or unlesioned hemisphere) for a given bird. P-values are from a two-sided Student’s t-test. **(D)** Integration of cell type specificity scores and ipsilateral versus contralateral coefficients to identify cell type-associated transcriptional effects of LMAN lesions. For the top 50 differentially expressed genes, cell type specificity was multiplied by log fold-change between ipsilateral and contralateral RA. These values were then split by sign then summed within each cell type to yield a differential expression (DE)-cell type score. Gray bars indicate the distribution (1% to 99%) of DE-cell type specificity scores for 100 random sets of 50 genes. **(E)** RA cell type specificity scores for top differentially expressed genes, divided into upregulated and downregulated genes. Values are scaled for each gene such that the cell type with the highest specificity score equals 1 and that with the lowest equals 0. At the top of each specificity score heatmap is the log fold-change expression for ipsilateral versus contralateral RA. **(F)** Distributions of Song DKL high vs. low log fold-change for the top 25 marker genes for each cell type. Dashed lines indicate the 1% and 99% median log fold-change for 100 randomly selected sets of 25 genes.

For each brain region, we performed comparisons between the region ipsilateral to the LMAN lesion to that in the contralateral hemisphere (Fig. 7C and Supplemental Table 6). As expected, RA exhibited the greatest expression changes between sides ipsilateral and contralateral to the lesion (35 genes with reduced and 40 genes with increased expression in ipsilateral, adjusted p-value < 0.1) compared to ipsilateral to contralateral comparisons of non-direct targets of LMAN (Fig. 7C,D). Genes that were more highly expressed ipsilateral to the lesion were enriched for immune responsive genes likely reflecting an injury response in RA to the afferent lesion (Fig. 7D,E). In contrast, genes with reduced expression ipsilateral to the lesion were enriched for a range of biological processes, including activity-dependent delayed primary response genes (Tyssowski et al., 2018), neuron cellular homeostasis, metalloendopeptidase activity, and potassium channels (Fig. 7E).

To examine more broadly how these expression alterations compared to those associated with deafening-induced song destabilization, we calculated the average ipsilateral versus contralateral fold change for the destabilization-associated gene modules described above (Fig. 7F). If LMAN lesions impose a molecular state associated with low variability and low plasticity, we would expect to see an inverse pattern of expression between Song D_KL_ and lesion differential expression. Indeed, on the whole, modules that had increased expression in RA with higher Song D_KL_ had lower expression ipsilateral to the lesion, and vice versa (Fig. 7G). However, one module, M74, showed an opposite pattern — it was the most strongly reduced module both with increased song destabilization and with LMAN lesions. M74 hub genes *CRHBP* and *SST* were reduced specifically in RA ipsilateral to the lesion and showed no change in other assayed regions (Fig. 7H and Figure 7-figure supplement 2A-C). This module is enriched for secreted neuropeptides, and the similarity of its expression change across both deafening and LMAN lesions could reflect its sensitivity to altered neural activity in RA, either through the loss of auditory input or through the direct loss of a major afferent to RA.

We integrated ipsi-vs-contralateral lesion differential expression with cell type specificity, as described above in the deafening analysis, to examine the cellular expression biases of genes that are influenced by the loss of LMAN. Upregulated genes were primarily expressed in non-neuronal cells and in particular in microglia and astrocytes, consistent with an injury response (Figure 7-figure supplement 2D,E). Indeed, marker genes for microglia show a strong increase in expression in RA ipsilateral to LMAN lesions, suggesting an increase in microglia abundance in RA (Figure 7-figure supplement 2F). This bias is consistent with a glial injury response to the lesion or alternatively may reflect glia-mediated synaptic plasticity. In contrast, downregulated genes were largely expressed in neurons (Fig. 7-figure supplement 2D,E), namely glutamatergic projection neurons (Glut-1) and MGE-derived GABAergic interneurons such as *Sst-*class (GABA-2), *Pvalb*-class (GABA-4), and cholinergic neurons (GABA-8).

## Discussion

Sensory feedback is necessary for the reliable and successful execution of learned motor skills, and its loss can lead to increased motor errors and aberrant motor plasticity. The deprivation of sensory experience has been used effectively to characterize plasticity within sensory systems and its underlying cellular and molecular mechanisms. In contrast, how sensory deprivation drives plasticity in associated sensorimotor circuitry at cellular and molecular levels is comparatively poorly understood. Here, we used the experimental advantages of birdsong, a highly precise learned motor skill that has a dedicated neural circuitry, to identify molecular pathways in sensorimotor circuits that are influenced by the loss of auditory input and associated vocal motor destabilization.

This model has particular relevance for understanding the neural basis of speech alterations caused by deafening that occurs after speech acquisition (post-lingual deafening). Similar to the effects of auditory feedback loss to birdsong, post-lingual deafening in humans reduces the rendition-to-rendition precision of speech production (Lane and Webster, 1991; Waldstein, 1990) and alters spectrotemporal features of speech (Lane and Webster, 1991; Schenk et al., 2003). Finally, the deterioration of both speech and birdsong is more extreme when deafening occurs at earlier ages, suggesting that there are similar age-dependent mechanisms of vocal stabilization in both systems (Brainard and Doupe, 2001; Cowie et al., 1982; Lombardino and Nottebohm, 2000; Waldstein, 1990). It is an open question how the different components of speech production neural circuitry respond to hearing loss at various biological levels, from molecular to physiological.

For the songbird, prior studies have identified a variety of circuit, cellular, and molecular mechanisms that may contribute to deafening-induced song-destabilization (Brainard and Doupe, 2000; Kojima et al., 2013; Mandelblat-Cerf et al., 2014; Peng et al., 2013, 2012a, 2012b; Scott et al., 2000; Tschida and Mooney, 2012; Wang et al., 1999; Watanabe et al., 2002; Zhou et al., 2017). These prior demonstrations, which have focused on a disparate set of song control structures and specific candidate mechanisms, motivated our interest in applying a circuit-wide and unbiased approach in this system to identify molecular responses to auditory deprivation-induced motor destabilization. Understanding these responses in the songbird vocal control system could provide insight into the neural mechanisms underlying the plasticity and resilience of both learned vocalizations and other well-learned motor skills.

### Molecular localization of song destabilizationanalysis

Past work on the neural mechanisms underlying song plasticity has largely focused on changes occurring in one or two brain regions at a time. Song destabilization in adult songbirds is associated with a variety of changes to the morphology and physiology of neurons in the song system including changes to dendritic spine stability and synapse densities in HVC and RA (Tschida and Mooney, 2012; Zhou et al., 2017); alterations to song tuning responses in LMAN (Roy and Mooney, 2007); and decreased synaptic inputs onto and increased intrinsic excitability of HVC projection neurons (Hamaguchi et al., 2014; Tschida and Mooney, 2012). Our neural circuit-wide analysis of gene expression responses to deafening allowed us to investigate which regions of the song system show the strongest transcriptional changes during song destabilization, providing a readout of the molecular correlates of neural plasticity. We found that the motor output nucleus RA showed the highest differential expression upon song destabilization, with substantial changes also found in Area X and to a lesser extent in HVC and LMAN. RA lies at the nexus of the motor pathway and the anterior forebrain pathway, a circuit required for song plasticity, and is a major locus of neural plasticity during juvenile song learning and adult song adaptation (Garst-Orozco et al., 2014; Miller et al., 2017; Ölveczky et al., 2011). This position in the song neural circuit makes it well situated to integrate neural activity associated with the stable motor program with AFP-generated contributions to deafening-induced plasticity.

Some of the most salient pathways upregulated in RA were associated with synaptic transmission and neuron spines, consistent with previous reports that found that deafening increases synapse densities, spine densities, and spine lengths in RA (Peng et al., 2012a; Zhou et al., 2017). These terms were also enriched to some extent for differentially expressed genes in HVC, in which neurons projecting to Area X exhibit decreased spine stability following deafening (Tschida and Mooney, 2012). Past work on several of the top differentially expressed genes in RA support a general role for altered synapse and spine dynamics during deafening-induced song plasticity. In particular, stathmin 1 (*STMN1*) is located at synapses and binds to tubulin to inhibit microtubule formation (Curmi et al., 1997; Shumyatsky et al., 2005). Knockout of *STMN1* in mice results in impaired long term potentiation in the amygdala and reduced memory in fear-conditioning tasks (Shumyatsky et al., 2005). Furthermore, *STMN1* is differentially phosphorylated during fear conditioning, altering its activity and AMPA receptor localization to the synapse (Uchida et al., 2014). Similarly, the surface glycoprotein *CD24*, which is upregulated in RA, influences neurite outgrowth (Gilliam et al., 2017) as well as synapse formation and transmission (Jevsek et al., 2006). Lastly, the lipid processing enzyme lipoprotein lipase (*LPL*) is upregulated in RA during song destabilization. Knockouts of *LPL* in mice result in impaired learning and memory, decreased presynaptic vesicles in the hippocampus (Xian et al., 2009), and reduced AMPA receptor expression (Yu et al., 2015).

The expression of neuropeptides was also broadly reduced following deafening across multiple song nuclei. This result suggests that song plasticity is a product of not only alterations to synapse structure and neurotransmitter-mediated signaling but also changes in neuromodulation. It has not been well-examined how secreted neuropeptides influence birdsong plasticity and neural activity in birdsong neural circuitry. However, extensive evidence garnered in other systems indicates that neuropeptide signaling has a powerful effect on neural circuit activity, plasticity, and behavioral output (Bargmann, 2012; Marder, 2011). The specific signaling systems altered following deafening in this study provide a set of candidate mechanisms that may influence song. For example, corticotropin releasing hormone binding protein (*CRHBP*), one of the most strongly downregulated genes in RA following deafening, modulates activity in the CRH signaling pathway (Kemp et al., 1998), which has diverse effects on long-term potentiation, neuronal excitability, and spine dynamics in central circuits (Aldenhoff et al., 1983; Blank et al., 2003; Chen et al., 2008; Fox and Gruol, 1993; Kratzer et al., 2013; Li et al., 2016). Such evidence suggests that the dynamic modulation of neuropeptides could play a prominent role in regulating birdsong stability and plasticity and may similarly influence the control of other stable sensorimotor skills such as human speech.

### Neuronal contributors to song plasticity

By integrating song system-wide and cell-resolved expression profiles, we can make initial predictions about which cell classes exhibit transcriptional changes during song destabilization. Glutamatergic projection neurons in RA are similar to layer 5 extratelencephalic neurons in the mammalian neocortex, both in terms of their projections to subcerebral structures and their expression profiles (Colquitt et al., 2021; Nevue et al., 2020; Pfenning et al., 2014; Vicario, 1991). A number of differentially expressed genes showed biased expression toward glutamatergic neurons in RA, including protein kinase C β (*PRKCB*), a calcium sensor associated with short-term plasticity (Chu et al., 2014; Fioravante et al., 2014) and previously shown to be upregulated in RA following deafening (Watanabe et al., 2002), as well as the lipid processing enzyme *LPL*, discussed above.

Our results also point to a prominent role for GABAergic interneurons in deafening-induced song plasticity. In particular, genes that had reduced expression with song destabilization showed an expression bias toward *Sst*- and *Pvalb-*class interneurons in RA. The interneuron subclasses present in the song system are strongly similar to well-characterized interneuron types in the mammalian neocortex, suggesting deep conservation of inhibitory networks (Colquitt et al., 2021). How these specific subclasses influence network activity in the song system is an open question; however, previous work has established a general role for local inhibition in the regulation of song learning and stability (Kosche et al., 2015; Vallentin et al., 2016). In particular, song learning in juveniles — during which song becomes more structured and less variable — refines the synaptic connectivity between glutamatergic projection neurons in RA and an inhibitory neuron type that has electrophysiological properties similar to fast-spiking *Pvalb*-class interneurons (Miller et al., 2017). Similarly, inhibitory input to HVC projection neurons increases and becomes more precise as song performance improves during juvenile song learning (Vallentin et al., 2016). Many of the *Sst/Pvalb*-biased genes affected by deafening-induced song plasticity are secreted neuropeptides that are sensitive to levels of neural activity (Hou and Yu, 2013; Tyssowski et al., 2018), suggesting that their reduced expression reflects reduced activity in these populations during birdsong destabilization. Moreover, several of these neuropeptides, including *SST* and *CRHBP*, act to inhibit neural activity, either directly through receptor binding or indirectly through interactions with other neuromodulators (Hou and Yu, 2013; Li et al., 2016; Pittman and Siggins, 1981).

This role for inhibition in maintaining birdsong structure has parallels to the role of inhibition during neural plasticity in mammals. For instance, the density of synapses from *Sst*- and *Pvalb-*class interneurons onto pyramidal neurons in the mammalian motor cortex is modulated during motor learning in mice, with an overall reduction of *Sst*-class input during motor plasticity (Chen et al., 2015). Likewise, low *Pvalb*-class network activity in the hippocampus is associated with increased synaptic plasticity, and low *Pvalb* expression, itself sensitive to neural activity, is found in the motor cortex during early motor learning (Donato et al., 2013). Similarly, increased neuronal excitability and decreased inhibition have also been found in the mammalian auditory cortex following deafening or noise trauma (Kotak et al., 2008, 2005; Seki and Eggermont, 2003). Together, these results suggest that reduced inhibition, either through altered synaptic transmission or neuromodulation, is a key component of neural plasticity in both the mammalian and avian central nervous system.

### Circuit contributions to transcriptional state

By sampling gene expression across the different connected components of birdsong neural circuitry in individual birds, our study allowed us to examine the correlation of gene expression in one brain area with that in another. Regions with direct projections to each other, for instance HVC to RA and LMAN to RA, tended to have a higher number of genes with correlated expression across individuals than song or non-song regions that are not directly connected. Moreover, genes with correlated expression across the three pallial (cortical-like) song regions HVC, RA, and LMAN were enriched for activity-dependent genes and secreted neuropeptides. These results could reflect the presence of a shared molecular state across the song system, perhaps established by singing-related neural activity or a common response to a general hormonal factor reflecting some aspect of a given bird’s state (e.g. testosterone levels which may in turn be affected by sensory deprivation (Livingston et al., 2000)). Two results further support that these correlations reflect some aspect of shared activity across regions. First, deafened birds exhibited, on the whole, reduced gene expression correlation between song regions, suggesting that either the loss of auditory information or associated song destabilization disrupts inter-region coordination of gene expression. Second, lesions of LMAN altered gene expression specifically in RA, one of its primary efferents, but not other brain regions to which it does not directly project. This analysis highlights the importance of considering inter-regional influences in understanding the mechanistic basis of transcriptional responses across an integrated neural circuit.

These cross-regional patterns of gene expression relate to a central question of this study: how does the loss of sensory input influence gene expression in sensorimotor circuits? Our results suggest a model in which altered activity propagates through existing circuits, such that the state of one circuit component progressively modifies gene expression in its targets. Local mechanisms engaged within each region, such as synapse/spine remodeling and neuropeptidergic signaling implicated here, could then alter circuit connectivity and function, leading to behavioral plasticity. Identifying how neural activity influences gene expression across neural circuits and what specific molecular and cellular factors in turn shape circuit function will be instrumental to better understand the neural mechanisms that underlie sensorimotor stability and its impairment following sensory loss.

## Materials and Methods

### Animal care and use

All Bengalese finches were from our breeding colonies at UCSF or were purchased from approved vendors. Experiments were conducted in accordance with NIH and UCSF policies governing animal use and welfare.

### Song recording and preprocessing

Birds were individually housed in wire cages in sound isolation chambers. Song was recorded using Countryman Isomax microphones taped to the top of the wire cage. Microphones were connected to USB preamplifiers that were connected to a Linux workstation. Audio was recorded at a frame rate of 44,100 samples/second using a custom python script, and, to select for periods of singing, blocks of continuous sound with amplitudes above a manually set threshold were saved as 24-bit WAV files.

### Song autolabeling

The analysis of specific spectral features (e.g. fundamental frequency) was performed on syllables that were labeled using a supervised machine learning approach, called hybrid-vocal-classifier or hvc (Nicholson, 2021). For each bird, 20-50 songs were manually labeled using the Matlab software evsonganaly. Using hvc, a set of spectrotemporal features was computed for each syllable (e.g. duration, mean frequency, pitch goodness, and mean spectral flatness as defined in (Tachibana et al., 2014)). These features and the manually defined labels were provided to *hvc* to train a support vector machine (SVM) with radial basis function, with a grid search across parameters C and gamma to identify parameters with the highest classification accuracy. A set of models were then trained using these selected parameters and a random sample of training syllables, and model accuracy was tested on a held-out set of syllables. For each bird, the number of input syllables and parameters were adjusted until label accuracy reached 95-100%. The model with the highest accuracy was then used to predict labels on unlabeled songs. To select confidently labeled syllables, a prediction confidence score was calculated for each syllable as the entropy (sklearn.stats.entropy) of the classification probabilities resulting from SVM model prediction. Syllables with a prediction confidence score greater than 0.5 were retained.

### Song dimensionality reduction

To project syllable spectrotemporal structure into a reduced dimension space, we used an approach developed by Sainburg et al. (Sainburg et al., 2020) with code and example scripts obtained from https://github.com/timsainb/AVGN. Songs were first isolated from audio recordings then segmented into syllables based on amplitude threshold crossings. Spectrograms were computed for each syllable using short-time Fourier transforms (512 window size, 0.5 ms step size, 6 ms window size, 44,100 frames per second) and frequencies between 500 Hz and 15,000 Hz were retained. Spectrograms were converted to mel scale using a mel filter with 128 channels. Syllables were compressed in the time dimension to a framerate of 640 frames per second then zero-padded to yield a standardized dimension of 128. Before dimensionality reduction, these 128×128 spectrograms were further reduced to 16×16 matrices then flattened yielding a 256 length feature vector for each syllable. Syllable x feature vector matrices were then processed using the single-cell analysis framework *Seurat v3 (Stuart et al., 2019)*. Principal component analysis was performed, then Uniform Manifold Approximation and Projection (UMAP) was performed on the first 10 principal components to produce a two-dimensional reduction.

### UMAP density differences

To calculate global differences in syllable spectral structure before and after a manipulation (e.g. deafening), we split each bird’s song UMAP by day relative to the manipulation and computed two-dimensional kernel density estimates (R package *MASS* v7.3 function *kde2d*, 200 x 200 grid) for each of the these per-day plots. A baseline UMAP structure was calculated as the mean density across the 2-4 days of singing before the manipulation, then density differences were calculated by subtracting this baseline density from each per-day density plot. Positive values from each difference plot were summed to give a single statistic for each day. Significance between hearing and deaf conditions for each day was determined using a two-sided t-test.

### Fundamental frequency statistics

To calculate fundamental frequency (FF) for a given harmonic stack, we first computed the average spectrogram for 20 randomly selected syllables. We then identified a time within the syllable (relative to syllable onset) with stable FF and defined minimum and maximum frequency bounds to define a frequency band containing the FF. A short-time Fourier transform (STFT) was then calculated at this time point using function *spec* from R package *seewave* v2.1.8 (Sueur et al., 2008) (1024 window size, 44,100 frames per second). FF was estimated by interpolating the frequency spectrum on an output vector spanning the minimum and maximum frequency bounds with a resolution of 1 Hz (function *aspline* from R package *akima* v0.6-2.2). The maximum value of this interpolated frequency spectrum was taken as the FF. Rolling variability of FF was calculated as the coefficient of variation (CV, standard deviation / mean) over a set of FF values for a given syllable and the 10 prior syllables (of the same type). To compare variability relative to a baseline period, FF CV values were transformed to a percentage relative to the average variability before manipulation. Group estimates and significance values were obtained from mixed effects linear models using R package *lme4* v1.1-27.1 *(Bates et al., 2015)* and function *lmer* (maximum likelihood criterion). The time period (before or after manipulation) was treated as a fixed effects and bird ID and syllable were treated as random effects with syllable nested under bird ID [model in *lme4* notation: period + (1 | bird / syllable)]. P-values for fixed effect were obtained using ANOVA (Type II, Wald chi-square test statistic, R package *car* v3.0-11 function *Anova,* (Fox and Weisberg, 2019)) followed by adjustments for multiple testing using Benjamini-Hochberg correction.

### Song D_KL_

To provide a single statistic that represents the amount of difference between songs in two conditions, we used a measure that we previously developed called Song D_KL_ (Mets and Brainard, 2018). Songs for a given bird were divided into ‘pre’ and ‘post’-procedure groups. The ‘pre’ group consisted of songs from at most four days before the procedure up to the day preceding the procedure. The ‘post’ group contained song from two days before the day of euthanasia to the day of euthanasia. A maximum of 50 songs were sampled from each day. Syllables were identified in song WAV files by amplitude thresholding using a manually defined threshold for each bird. Mean power spectral densities (PSD) were computed for each segmented syllable using short time Fourier transforms via R package seewave v2.1.8 (Sueur et al., 2008) and function meanspec (window length ‘wl’ = 512, overlap ‘ovlp’ = 0%, normalized ‘norm’ = T). Syllables in each dataset were split into a training dataset of 500 syllables and a held-out dataset of the remaining syllables. 50 PSDs were randomly selected from the ‘pre’ training dataset to serve as reference syllables for distance calculations. Inter-syllable spectral distances were calculated as Euclidean distances between this reference syllable set and each PSD, generating distance matrices for the ‘pre’ training and held-out datasets and the ‘post’ training and held-out datasets. Gaussian mixture models (GMMs) were fit to the ‘pre’ training distance matrix using function Mclust (5-12 mixture components, diagonal multivariate mixture model with varying volume, varying shape “VVI”) from R package mclust v5.4.7 (Scrucca et al., 2016). Bayesian Information Criterion (BIC) was computed for each model and second-order differences (difference of the difference) were calculated between the BICs for models with increasing numbers of mixture components. The model with minimum second-order BIC difference was selected for further use. A GMM was likewise fit to the ‘post’ training distance matrix using the same number of mixture components as in the selected ‘pre’ training model. The likelihood of generating each syllable in the ‘pre’ held-out dataset under the ‘pre’ and ‘post’ GMMs was calculated. This procedure was repeated ten times with different randomly selected reference syllables. The Kullback-Leibler divergence was then calculated as

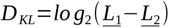

where 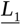 is the mean likelihood of observing a ‘pre’ held-out syllable across the ten replicated ‘pre’ models and 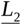 is the corresponding mean likelihood value for the ‘post’ models. These syllable-level *D_KL_* values were then averaged to give a single *Song D_KL_* for a given bird.

### Deafening by bilateral cochlea removal

Nine Adult male Bengalese finches (103-458 days post hatch, median±SD of 133±123) were deafened by bilateral cochlea removal. Birds were anesthetized using isoflurane and an incision was made in the skin covering the ear canal to expose the canal. The tympanic membrane was ruptured, and the columella removed using forceps. Cochlea were removed using a fine tungsten wire shaped into a hook. The incision was then resealed using VetBond (3M). For each deafened bird, a control (‘hearing’) bird underwent a sham surgery on the same day in which the bird was anesthetized and the skin incision was made then resealed. Birds survived for 4, 9, or 14 days (3 hearing and 3 deaf birds for each timepoint) then were euthanized as described in *Euthanasia and brain preparation*.

### Unilateral LMAN lesions

Five birds received unilateral LMAN lesions, three with left-hemisphere lesions and two with right-hemisphere lesions. LMAN was electrolytically lesioned using a 100 kOhm platinum/iridium electrode. LMAN was stereotactically located at 4.7 mm AP, 1.7 mm ML, and 2.1 mm DV using a beak angle of 50 degrees. In one hemisphere, five penetrations were made, one at the given coordinates and four more +/− 300 microns from this center position. At each penetration, 100 μA of current at the anode was passed for 60 seconds. At experiment end, birds were euthanized as described in *Euthanasia and brain preparation*. To assess lesion completeness, 20 um coronal cryosections were collected at 100 um intervals across the anterior-posterior extent of LMAN onto SuperFrost Plus slides (FisherBrand), then Nissl stained as described in *Standard Nissl stain, fresh-frozen cryosections*. The volume of LMAN was estimated in ImageJ/Fiji (Schindelin et al., 2012) by calculating the area of LMAN on the unlesioned side in each section that it was visible, interpolating a smooth curve in R across these measured areas and the known distance between cryosections, then calculating the area under the curve. This procedure was repeated for any residual LMAN visible on the lesioned side, and lesion percentage was calculated as 100 * (1 − [volume LMAN lesioned] / [volume LMAN unlesioned]). LMAN was considered lesioned if more than 25% of volume was spanned by the lesion.

#### Standard Nissl stain, fresh-frozen cryosections

Frozen sections were allowed to come to room temperature for at least 20 minutes then placed in a glass staining rack. Then slides were sequentially transferred to two rounds of xylenes for 5 minutes, two rounds of 100% ethanol for 5 minutes, one round of 95% ethanol for 5 minutes, 1 round of 70% ethanol for 5 minutes, water for 1 minute, stained in 0.5% cresyl violet solution for 30 minutes, then rinsed for 1 minute in water. Slides were then transferred to one round of 70% ethanol for 15-20 seconds (depending on desired staining intensity), one round of 95% ethanol for 30 seconds, two rounds of 100% ethanol for 30 seconds each, then two rounds of xylenes for 3 minutes each. DPX Mountant (Sigma) was applied, then slides were coverslipped. 0.5% cresyl violet was prepared as 300 mL water, 1 mL glacial acetic acid, and 1.5 g cresyl violet acetate. Solution was stirred for two days with no heat then filtered.

### Euthanasia and brain preparation

Birds were euthanized using isoflurane, decapitated, and debrained. All birds used for Serial Laser Capture RNA-seq were euthanized two hours after lights on at 9 AM. Brains were flash-frozen in −70 C dry ice-chilled isopentane for 12 seconds within 4 minutes from decapitation.

### Serial laser capture microdissection RNA-sequencing (SLCR-seq) — overview

We were motivated by improvements to low-input RNA-sequencing stemming from optimized single-cell approaches to develop a method that would allow the construction of tens to hundreds of gene expression libraries from anatomically-defined regions. To achieve this we combined an optimized rapid Nissl staining protocol, laser capture microdissection, scalable RNA purification, and low-cost and low-input RNA-sequencing library construction into a single pipeline called Serial Laser Capture Microdissection RNA-sequencing (SLCR-seq).

#### SLCR-seq — cryosectioning

Surfaces in the cryostat chamber were first cleaned using a mixture of 50% RNaseZap (Ambion)/50% ethanol followed by a rinse of 70% ethanol in nuclease-free water. Flash-frozen brains were removed from −80 °C storage and allowed to equilibrate in a cryostat chamber set to −18 °C for ~30 minutes. PEN membrane slides for LCM (Leica) and Superfrost Plus glass slides for histology (Fisherbrand) were placed in the cryochamber to chill. Once equilibrated, the brain was mounted onto a cryostat chuck using a small amount of OCT (TissueTek) with the posterior surface down and the anterior surface available for coronal sectioning. The brain was trimmed approximately 1.8 mm until reaching the anterior-posterior position of LMAN and Area X, which were visible as slightly darker regions. Sections were cut at 20 μm, transferred to pre-chilled membrane or glass slides, then melted onto the slides using a metal dowel that was pre-warmed on a slide warmer. Once a section was fully melted, the slide was transferred to a metal block in the cryostat chamber to refreeze the section. After sectioning through LMAN and Area X, the brain was detached from the chuck and remounted along the cut anterior surface for sectioning from the posterior surface. The brain was trimmed until reaching the anterior-position for RA (~0.8 mm from the posterior surface of the forebrain), which was also evident as a slightly darker region. Sections were collected onto membrane and glass slides as described through the level of HVC (~2.3 mm from posterior surface) or Field L (visible as a dark curve extending from the medial surface). Once collection was finished, slides were transferred to plastic slide mailers and stored in freezer boxes at −80 °C. Remaining brain tissue was re-wrapped in aluminum foil, placed back into a 15 mL conical tube, and stored at −80 °C. Test assays indicated that brains could be resectioned once more (for a total of two sectioning sessions) without negatively impacting RNA quality.

### SLCR-seq — rapid Nissl stain

A fast Nissl staining procedure was developed to quickly stain cryosections before laser capture microdissection. Anhydrous 100% ethanol solution was prepared by adding 15 g of molecular sieve beads (Sigma 208582, 3 Å, 8-12 mesh) to 500 mL 100% molecular grade ethanol (Sigma E7023). Cresyl violet staining solution was prepared as by dissolving cresyl violet powder (Sigma) to 4% wt/vol in 75% ethanol (75% molecular grade ethanol, 25% nuclease-free water), stirring for two days, then filtering through a 0.22 μm filter. Before staining, a series of 95%, 75%, 50% ethanol solutions were prepared. To stain, each slide was thawed at room temperature on a bench for 20 seconds then transferred to 95% ethanol for 30 seconds, 75% ethanol for 30 seconds, and 50% ethanol for 30 seconds. 400 μL of cresyl violet staining solution was then applied to the slide for 30 seconds. Slides were destained and dehydrated by transferring to 50% ethanol for 30 seconds, 75% ethanol for 30 seconds, 95% ethanol for 30 seconds, then two rounds of 100% ethanol for 30 seconds each. Slides were then allowed to air dry. Time series experiments indicated that RNA quality was maintained for up to 45 minutes following staining.

### SLCR-seq — Laser capture microdissection

After staining, slides were loaded onto a Leica LMD7000. Song nuclei were identified by anatomical landmarks (such as lamina and position relative to brain surfaces) and their higher intensity Nissl staining relative to surrounding regions. Sections cut from the surrounding tissue (power 45, aperture 50, speed 10, specimen balance 0, head 90%, pulse 92, offset 15) into 8-well strip caps containing 31.5 μL of RNA Lysis Buffer/PK (see SPRI RNA purification). After filling each cap with a section, the strip was placed onto a 96-well plate pre-chilled on ice and covered with an ice pack. Once a plate was filled, it was vortexed, spun down at 3,250 x g for 5 minutes at 4 °C, then transferred to dry ice. For long-term storage, plates were stored at −80 °C.

### SLCR-seq — SPRI RNA purification

The following solutions were prepared before LCM section collection: 50% guanidinium thiocyanate (Sigma) in nuclease-free water, 5X CN buffer (250 mM sodium citrate pH 7.0 (Sigma), 5% NP-40 (Sigma)), and RNA Lysis Buffer (20% guanidine thiocyanate, 1X CN buffer). The following solutions were prepared before RNA purification: RNA Wash Buffer (25 mM sodium citrate pH 7.0, 15% guanidinium thiocyanate, 40% isopropanol) and solid phase reversible immobilization (SPRI) bead solution. SPRI bead solution was prepared by first vortexing Sera-Mag SpeedBeads™ Carboxyl Magnetic Beads, hydrophobic (Fisher) until fully suspended transferring 1 mL beads to a 1.5 mL tube. Beads were washed by placing the tube on a tube magnet, waiting until solution cleared, removing the solution, adding 1 mL of TE Buffer (10 mM UltraPure Tris HCl, pH 8.0 (ThermoFisher), 1 mM EDTA pH 8 (ThermoFisher)), and pipetting to mix. This wash was repeated once more, then the beads were resuspended in 1 mL TE Buffer. Separately, 9 g polyethylene glycol 8000 (Amresco), 10 mL 5 M NaCl, 500 μL 1 M UltraPure Tris HCl pH 8.0 (ThermoFisher), 100 μL 0.5 M EDTA pH 8.0 (ThermoFisher), and 500 μL 2% sodium azide (Sigma) were combined and brought to ~49 mL using nuclease-free water. Solution was mixed by inversion until PEG 8000 went into solution. Then, 137.5 μL of 20% Tween-20 and 1 mL of beads/TE were added and mixed by inversion. This SPRI bead solution was then stored at 4 °C.

Just before LCM collection, 31.5 μL of RNA Lysis Buffer/PK (1.5 μL of Proteinase K (Ambion), 30 μL of RNA Lysis Buffer) was prepared for each well. To purify RNA following LCM section collection, samples were first allowed to thaw on ice if stored at −80 °C. SPRI bead solution was allowed to come to room temperature, then 40 uL SPRI bead solution was mixed with 47.5 uL isopropanol for each sample. Samples were then lysed by incubating at 42 °C for 30 minutes in a thermocycler then placed at room temperature. 87.5 uL of SPRI/isopropanol solution was added to each sample then mixed 10x by pipetting. Samples were incubated for 5 minutes at room temperature then transferred to a magnetic plate stand. After 3 minutes, the solution was removed, the plate was removed from the magnetic stand, 100 uL of RNA Wash Buffer was added, and beads were resuspended by pipetting. The plate was immediately transferred back to the magnetic plate stand and held there for 2 minutes until the solution cleared. The solution was removed, and the plate was removed from the stand. 100 uL of 70% ethanol was added, beads were resuspended by pipetting 10 times, the plate was returned to the magnetic stand, the solution was allowed to clear for 2 minutes, the solution was removed. This step was repeated for two total ethanol washes. Following the final wash, the beads were allowed to dry for 10 minutes while the plate remained on the stand. Residual ethanol was removed by pipetting. To elute RNA, the plate was removed from the magnet, 15 uL of nuclease-free water was added to each sample, and beads were resuspended by pipetting 10 times. Samples were incubated at room temperature for 5 minutes, then the plate was transferred to a low-elution volume magnetic stand. After 2 minutes or until the solution cleared, 10-12 μL eluted RNA was transferred to new 96-well plates on ice. Plates were sealed using foil adhesive, frozen on dry ice, then transferred to −80 °C for long-term storage.

### SLCR-seq — library preparation

The SLCR-seq library preparation was adapted from several low input and single-cell RNA-sequencing library protocols (Islam et al., 2014, 2012; Kivioja et al., 2012; Macosko et al., 2015; Picelli et al., 2014, 2013). Barcoded unique molecular identifier (UMI) reverse transcription (RT) primers were prepared in advance in a 96-well plate (RT/TSO/dNTP mix). Each well contained 10 μM barcoded reverse transcription primer (RT_primer, IDT), 10 μM template-switching oligonucleotide with lock nucleic acids (TSO_LNA, Exiqon), and 10 mM dNTPs. Plates were sealed with foil adhesive and stored at −80 °C. Two RT primers were used in this study: one for the initial 18 bird deafening dataset (RT_primer_v1, 25 base UMI, 6 base barcode), and another for the 10 bird unilateral LMAN dataset (RT_primer_v2, 14 base UMI, 12 base barcode). RT_primer_v1 and RT_primer_v2 sets consisted of 24 and 48 barcodes, respectively (Supplemental Table 1). Barcodes were at least one edit distance away from all other barcodes in the set.

For library preparation, total RNA prepared from *SPRI RNA purification* was thawed on ice, then 4 μL total RNA was placed into a well of a 96-well plate chilled on ice. 1 μL RT/TSO/dNTP mix was added and mixed 10 times by pipetting. Plates were sealed with foil adhesive, incubated at 72 °C for 3 minutes, then snap-cooled in ice for at least 2 minutes. An RT Master Mix was prepared containing 1x Enzscript RT buffer (Enzymatics), 5 mM dithiothreitol, 1 mM betaine, 12 mM MgCl_2_, 0.25 μL Recombinant Ribonuclease Inhibitor (Takara), and 10 U/μL Enzscript Moloney-Murine Leukemia Virus Reverse Transcriptase (Enzymatics). 5 μL of RT Master Mix was added to each sample and mixed by pipetting 10 times. Plates were sealed with foil adhesive and incubated in a thermocycler: 42 °C for 90 minutes, 70 °C for 15 minutes, 4 °C hold. Reactions were then pooled within a barcode set (e.g. barcodes 1-48 from RT_primer_v2 were combined into one tube). To purify cDNA, 0.6x volume of Ampure XP bead solution was added to each pooled sample and mixed by pipetting 10 times. Samples were incubated for 5 minutes and transferred to a tube magnet. After the beads cleared from the solution, the solution was removed, and the beads were washed in 400 μL freshly prepared 80% ethanol for 30 seconds. This step was repeated for a total of two washes. After the second wash, the ethanol solution was removed, and the beads were allowed to dry for 5-10 minutes. Beads were then resuspended in 22 μL of nuclease-free water and incubated for 2 minutes. 20 μL eluted cDNA was transferred to new 1.5 mL LoBind tubes or 96-well plates and either stored at −20 °C or amplified immediately.

During the purification a 40 μL cDNA Amplification Master Mix was prepared containing 10 μL KAPA HiFi 5x Buffer, 1 μL 10 mM dNTPs, 4 μL 10 mM TSO_PCR primer, 0.5 μL 1 U/μL KAPA HiFi Hotstart DNA polymerase, and 24.5 μL nuclease-free water. 10 μL of purified cDNA was added to this master mix, pipetted 10x to mix, then amplified under the following cycling parameters: 95 °C for minutes, then 4 cycles of 98 °C for 30 seconds, 65 °C for 45 seconds, and 72 °C for 3 minutes. Reactions were then placed on ice. During this initial amplification, a second master mix was prepared to determine the target number of amplification cycles by quantitative PCR. This mix contained 3 μL KAPA HiFi 5x Buffer, 0.3 μL 10 mM dNTPs, 1.2 μL 10 mM TSO_PCR primer, 0.15 μL 1 U/μL KAPA HiFi Hotstart DNA polymerase, 0.75 μL 20x EvaGreen (Biotium), and 4.6 μL nuclease-free water. 5 μL of preamplified cDNA was added to this mix and amplified in a real-time PCR machine: 98 °C for 3 minutes, followed by 24 cycles of 98 °C for 20 seconds, 67 °C for 20 seconds, and 72 °C for 3 minutes, followed by 72 °C for 5 minutes. The target number of additional cycles was determined by identifying the Ct at 20% of the max fluorescence then subtracting 5 cycles from this number. This number was generally between 5-7 additional cycles. The remaining 45 μL was placed back into the thermocycler and cycled for 98 °C for 30 seconds, the number of additional cycles at 98 °C for 20 seconds, 67 °C for 20 seconds, and 72 °C for 3 minutes, followed by 72 °C for 5 minutes.

To purify the amplified cDNA, 0.6x volume of Ampure XP bead solution was added to each reaction and mixed by pipetting 10 times. Samples were incubated for 5 minutes and transferred to a tube magnet. After the beads cleared from the solution, the solution was removed, and the beads were washed in 200 μL freshly prepared 80% ethanol for 30 seconds. This step was repeated for a total of two washes. After the second wash, the ethanol solution was removed, and the beads were allowed to dry for 5 minutes. Beads were then resuspended in 22 μL of nuclease-free water and incubated for 2 minutes. 20 μL eluted cDNA was transferred to new 1.5 mL LoBind tubes or 96-well plates and stored at −20 °C. Sample concentration was quantified using Qubit dSDNA High Sensitivity kit (ThermoFisher), then sample concentrations were standardized to 100 pg/μL.

To prepare tagmented DNA, 4 μL (400 pg) of amplified cDNA was added to 10 μL Tagmentation Buffer (Buffer TD from the Nextera XT DNA Sample Prep Kit, Illumina), 1 μL nuclease-free water, and 5 μL ATM (Nextera XT). Reactions were mixed by pipetting 10 times the incubated at 55 °C for 5 minutes. 5 μL of Buffer NT was then added, then the reactions were incubated for 5 minutes at room temperature.

Final libraries were constructed by first preparing a PCR master mix containing 20 μL KAPA HiFi 5x Buffer, 2 μL 10 mM dNTPs, 5 μL 10 mM P5-TSO_Hybrid primer, 5 μL 10 mM PCR2 primer, 1 μL 1 U/μL KAPA HiFi Hotstart DNA polymerase, and 42 μL nuclease-free water. PCR2 contains an i7 index (Supplemental Table 1). The 25 μL tagmentation reaction was then added directly to the mix, and mixed by pipetting 10 times. Samples were amplified using 72 °C for 3 minutes; 95 °C for 3 minutes; followed by 16 cycles of 98 °C for 10 seconds, 55 °C for 30 seconds, and 72 °C for 30 seconds; followed by 72 °C for 5 minutes. Samples were then purified by adding 1.2x volumes of Ampure XP, incubating for 5 minutes, then transferring to a tube magnet. After the beads cleared from the solution, the solution was removed, and the beads were washed in 200 μL freshly prepared 80% ethanol for 30 seconds. This step was repeated for a total of two washes. After the second wash, the ethanol solution was removed, and the beads were allowed to dry for 5 minutes. Beads were then resuspended in 22 μL of Low Elution Buffer (10 mM Tris HCl pH 8.0, 0.1 mM EDTA, 0.05% Tween-20) and incubated for 2 minutes. 20 μL eluted cDNA was transferred to new 1.5 mL LoBind tubes and stored at −20 °C. Library size distributions were assessed using a Bioanalyzer High Sensitivity DNA Chip (Agilent), and library concentrations were determined using the KAPA Library Quantification Kit (Illumina Complete Kit, Roche). Samples were pooled at equal concentrations then size selected using a BluePippin and 2% BluePippin gels. DNA from 180 to 500 bp was selected then purified using the MinElute kit (Qiagen) with two rounds of 10 μL elution in Low Elution Buffer. Samples were stored at −20 °C.

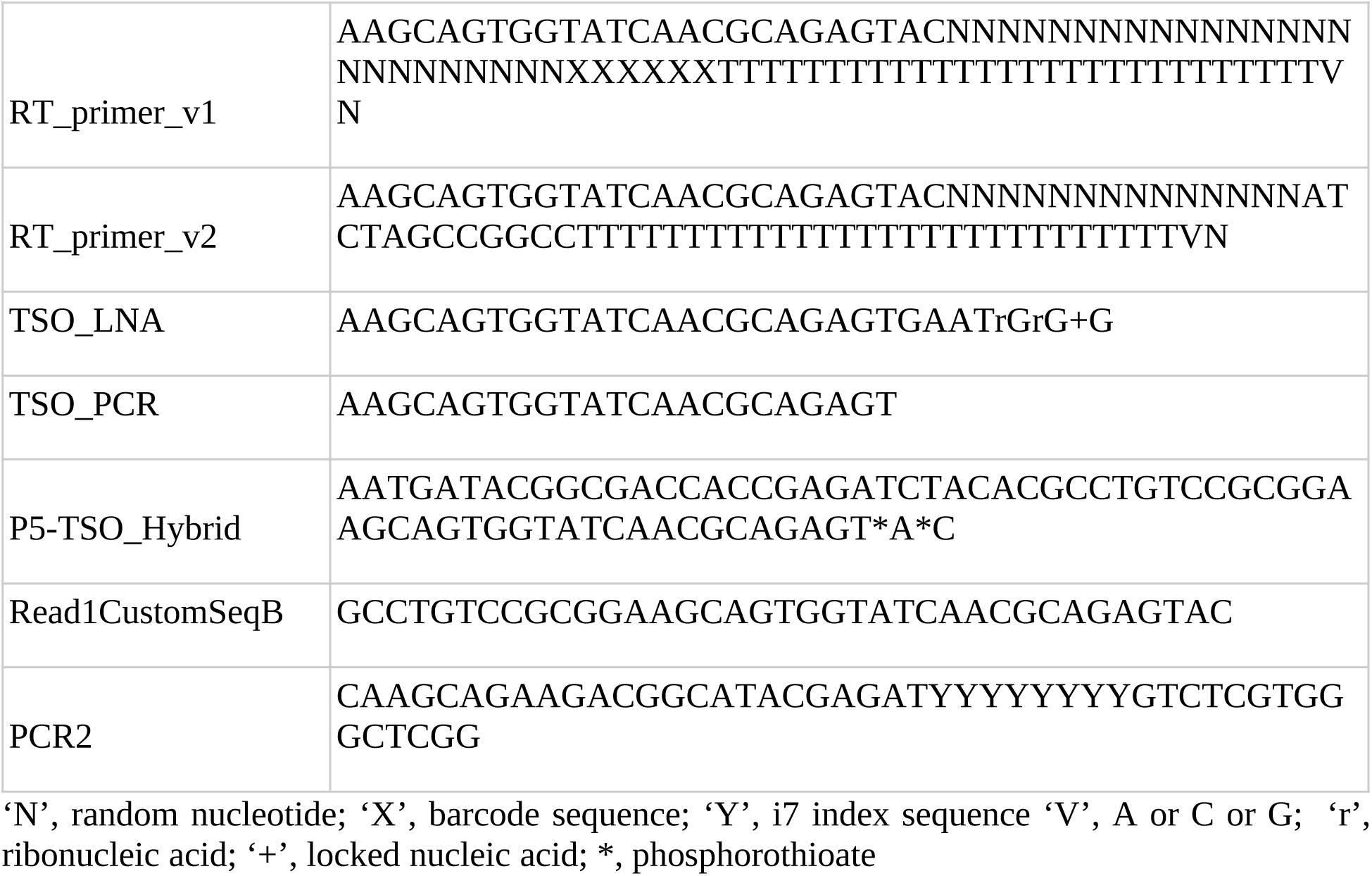

### RNA-sequencing preprocessing

Sequencing reads were first trimmed for adaptor sequences using trim_galore (https://github.com/FelixKrueger/TrimGalore, --quality 20, --paired, --overlap 10, adaptors AAAAAAAAAA and GTACTCTGCGTTGATACCACTGCTTCCGCGGACAGGCGTGTAGATCT). We first generated an initial alignment to the Bengalese finch genome (lonStrDom2, GCF_005870125.1) using *STAR* v2.7.8a (STARsolo mode, default parameters, --outFilterIntronMotifs RemoveNoncanonical) (Dobin et al., 2013). To better annotate the 3’ UTRs of Bengalese finch gene models, we identified transcript 3’ ends by assembling transcripts using these initial alignments and the RNA-seq assembler *Stringtie* (Kovaka et al., 2019) (--fr -m 100). These Stringtie models were then intersected with the NCBI Bengalese finch transcriptome (lonStrDom2, GCF_005870125.1). New Stringtie exons were filtered by same-strandedness to the intersected reference genes, minimal expression level (at least 10% of expression max for a given gene), and at least within 10 kilobases from the 3’ end of the gene. 3’UTRs of the reference transcriptome were extended out to these new exons. Reads were then re-aligned to this extended transcriptome using the ‘bus’ subcommand from *kallisto* v0.46.1 (Bray et al., 2016; Melsted et al., 2021) followed by barcode error correction using *bustools* v0.39.3 ‘correct’, sorting using ‘sort’, and read counting using ‘count’.

### Differential expression analysis

Gene-sample count matrices were filtered to remove lowly expressed genes, defined as having a total number of reads across samples less than the number of samples divided by eight (the number of brain regions assayed). For each sample we also calculated the ‘cellular detection rate’ (CDR) or the number of genes detected in a given sample, previously shown to substantially influence differential expression analysis on single-cell RNA-sequencing samples (Finak et al., 2015). Low quality samples were defined as having a CDR less than 30% of the total number of genes in the reference annotation (18,674 genes). Normalization factors were calculated using the function *calcNormFactors* from the R package *edgeR* v3.31.4 and the “TMMwsp” method. The count matrix, these normalization factors, and a design matrix were then provided to the function *voom* from the *limma* package v.3.48.3 (Law et al., 2014; Ritchie et al., 2015). The design matrix was specified as:

~0 + position + position:num_songs_on_euth_date_log_scale + position:kl_mean_log_scale_cut2_proc2 + position:kl_mean_log_scale_cut2_proc2:num_songs_on_euth_date_log_scale + position:nsongs_per_day_pre_log_scale + cdr_scale + frac_mito_scale + sv1 + sv2

where ‘position’ is an indicator for brain region, ‘num_songs_on_euth_date_log_scale’ is log-transformed total number of songs sung on the day of euthanasia, ‘kl_mean_log_scale_cut2_proc2’ is log-transformed Song D_KL_ discretized into three equally sized bins, ‘nsongs_per_day_pre_log_scale’ is log-transformed average number of song sung per day during the pre-procedure period, ‘cdr_scale’ is CDR, and ‘frac_mito’ is the fraction of reads mapping to mitochondrial genes in a given sample. Variables with ‘scale’ in their names were mean-subtracted and standard deviation-normalized. ‘sv1’ and ‘sv2’ correspond to the top two surrogate variables calculated using the function *svaseq* from the R package *sva* v3.40.0 (Leek, 2014), with full model specified as above and null model given as “~0 + position + cdr_scaled + frac_mito_scale”. To calculate the within-bird correlation between samples, the resulting voom object was passed to *duplicateCorrelation* with block specified as the bird ID (for the deafening samples) or bird ID and brain hemisphere (for the unilateral LMAN lesion samples). To fit the above model, the voom object, design matrix, and the consensus within-bird correlation were input to function *lmFit* from limma. Coefficient estimates and standard errors for each coefficient were calculated using function *contrasts.fit*, the function *eBayes* was used to compute moderated t-statistics and p-values, and the function *topTable* was used to adjust p-values using the Benjamini-Hochberg method. Genes were considered differentially expressed if their adjusted p-values were less than 0.1.

Differentially expressed genes in the unliateral LMAN lesion SLCR-seq dataset was calculated similarily but with design specified as:

~0 + group + tags + cdr_scale

where group indicates whether the region is ipsilateral or contralateral to the LMAN lesion. The variable ‘tags’ refers to bird ID tag and therefore controls for bird-level differences allowing pairwise comparisons of the effect of lesioning within birds.

Expression estimates and standard errors for a given bird and brain region were computed using a regression approach with a design matrix specified as:

~0 + position:tags + cdr_scale + frac_mito_scale

where ‘position’ is an indicator for brain region, ‘tags’ is the bird ID, ‘index2’ is a categorical variable indicating the sequencing run, and ‘cdr_scale’ and ‘frac_mito_scale’ are as described above. Standard errors were extracted from the linear fit model ‘fit’ as: sqrt(fit$s2.post) * fit$stdev.unscaled.

### Network analysis

We used the R package *MEGENA* (Multiscale Clustering of Geometrical Network, v1.3.7) to identify modules of genes with correlated expression across SLCR-seq data (Song and Zhang, 2015). Low quality samples were removed by retaining samples with cellular detection rates above 0.42. Samples expression values were normalized using normalization factors calculated as above (*calcNormFactors* and the “TMMwsp” method) then log-transformed with a pseudocount of 1. Samples were then split by brain region. To remove batch effects contributed by which pool a given sample was in, we used the function *ComBat (Johnson et al., 2007)* from the R package *sva* v3.40.0. For each brain region, we then selected the top 2000 variable genes as defined using the *Seurat* function *FindVariableFeatures* v4.0.4 and the ‘vst’ method. Signed Pearson correlations between every pair of genes were then calculated using the function *calculate.correlation* from *MEGENA,* which calculates false discovery rates by permutation (50 permutations). Correlations with an FDR less than 0.05 were retained. We passed these pairwise correlations to function *calculate.PFN* to generate a more sparse network that retains information edges using the MEGENA Planar Filtered Network algorithm. Module detection was then performed on this filtered network using the function *do.MEGENA*.

To identify modules associated with behavioral features, we calculated the average of log-transformed estimates for a given coefficient across genes in a given module. To identify modules with greater (or lesser) than expected fold-changes, for each module we randomly selected the same number of genes and averaged their log-transformed coefficient estimates 100 times. Modules that had averages less than 1% or greater than 99% of this null distribution were considered significant. Hub genes were designated using the approach defined in MEGENA. For each module, the link weights of the planar filtered network were permuted 100 times to generate a set of random networks. Within-module connectivities, defined as the sum of link weights with each other gene in a gene’s module, were calculated for each gene in each random network. The p-value was calculated as the probability of finding within-module connectivity values from this null distribution equal to or greater than the observed within-module connectivity. These p-values were then adjusted using the Benjamini-Hochberg method and genes with adjusted p-values less than 0.05 were designated hub genes.

To compute gene module memberships, eigengenes were first determined for each module using the R package *WGCNA* v1.70-3 (Langfelder and Horvath, 2008) and function *moduleEigengenes* then the Pearson correlation was computed between each module eigengene and each gene. Module preservation statistics were calculated using the *WGCNA* function *modulePreservation (Langfelder et al., 2011)*.

### Gene set enrichment analysis

Gene Ontology lists were obtained from the Molecular Signatures Database (set C5, version 7). Gene set enrichment analysis was performed using the R package *fgsea* v1.18.0 (Korotkevich et al., 2021). T-statistics from *voom* regression or gene module membership scores from *MEGENA* were input into the function *fgseaMultilevel* (minSize = 20, maxSize = 200). Resulting pathways were filtered for those with an adjusted p-value less than 0.2 and similar pathways were pruned using *collapsedPathways*(pval.threshold = 0.01 or 0.05).

### Inter-region correlation analysis

To analyze inter-region gene correlations, we first selected in each region the 500 genes with the highest variability, computed as the variance-mean ratio of non-log expression across samples. For each gene, we calculated Pearson correlation values for each pairwise combination of brain regions, yielding region-by-region correlation matrices. To generate null distributions for each gene, we shuffled bird identities for each pairwise region-region comparison 100 times and computed Pearson correlations. We thresholded observed correlations using statistics from these shuffled distributions (correlation lesser or greater than the 2.5% or 97.5% shuffled quantiles). We calculated the across-bird expression similarity between regions as the number of thresholded correlations.

To determine if deafening alters inter-region gene expression coupling in the song system, we computed pairwise Pearson correlations for each gene between each region for hearing and deaf birds separately, then took the absolute value for each matrix. The hearing absolute correlation matrix was then subtracted from the deaf absolute correlation matrix. This procedure was repeated on 100 shuffled distribution to generate a null distribution of differential absolute correlations. Differentially correlated genes were called those with an deaf versus hearing value less than (decorrelation) or greater than (correlation gain) extreme values of a shuffled distribution calculated for each pairwise comparison (2.5% or 97.5%, respectively).

To determine if deafening alters inter-region gene expression coupling in the song system, we computed pairwise Pearson correlations for each gene between each region for hearing and deaf birds separately, then took the absolute value for each matrix. The hearing absolute correlation matrix was then subtracted from the deaf absolute correlation matrix. This procedure was repeated on 100 shuffled deaf and hearing matrices. Differentially correlated genes were called those with an deaf versus hearing value less than (decorrelation) or greater than (correlation gain) extreme values of a shuffled distribution calculated for each pairwise comparison (2.5% or 97.5%, respectively)

### Cell type specificity and differential expression scores

For each gene, a specificity score was calculated as 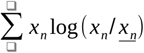, where *x_n_* is expression divided by the sum of expression across all clusters and 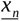 is the mean of this value. Regression coefficient-cell type specificity scores were calculated by selecting differentially expressed genes (adjusted p-value < 0.1) then splitting genes by the sign of the coefficient. Scores were then computed as the dot-product between the cell type x gene specificity matrix and the gene x coefficient matrix.

### Fluorescent in situ hybridization (FISH)

FISH was performed using the hairpin chain reaction system from Molecular Instruments. Birds were euthanized using isoflurane, decapitated, and debrained. Brains were flash-frozen in −70 °C dry ice-chilled isopentane for 12 seconds within 4 minutes from decapitation then stored at −80 °C. Fresh-frozen brains were cryosectioned at 16 μm onto SuperFrost slides (Fisherbrand) chilled in the cryochamber then melted onto the slide using a warmed metal dowel. Slides were then transferred to −80 °C for storage. For the FISH, slides were transferred from −80 °C to slide mailers containing cold 4% PFA and incubated for 15 minutes on ice. Slides were washed three times for 5 minutes using DEPC-treated PBS + 0.1% Tween-20, dehydrated in 50%, 70%, and two rounds of 100% ethanol for 3-5 minutes each round, then air dried. Slides were then transferred to a SlideMoat (Boekel Scientific) at 37 °C. 100 μL of v3 Hybridization Buffer (Molecular Instruments) was added to each slide, which were then coverslipped and incubated for 10 minutes at 37 °C. Meanwhile, 2 nM of each probe was added to 100 μL Hybridization Buffer and denatured at 37 °C. Pre-hybridization buffer was removed, 100 μL of probe/buffer was added, slides were coverslipped and incubated overnight at 37 °C. The next day, coverslips were floated off in Probe Wash Buffer (PWB, 50% formamide, 5x SSC, 9 mM citric acid pH 6.0, 0.1% Tween-20, 50 μg/ml heparin), then washed in 75% PWB/25% SSCT (5x SSC, 0.1% Tween-20), 50% PWB/50% SSCT, 25% PWB/75% SSCT, 100% SSCT for 15 minutes each at 37 °C. This was followed by 5 minutes at room temperature in SSCT. Slides were incubated in 200 μL of Amplification Buffer (provided by company) for 30 minutes at room temperature. Alexa fluor-conjugated DNA hairpins were denatured for 90 seconds at 95 °C then allowed to cool for at least 30 minutes in the dark at room temperature. Hairpins were added to 100 μL amplification buffer, applied to slides, and incubated overnight at room temperature. The following day, slides were washed in SSCT containing 1 ng/mL DAPI for 30 minutes at room temperature, then SSCT for 30 minutes at room temperature, followed by a final 5 minutes in SSCT at room temperature. Prolong Glass Antifade Medium (Thermofisher) was added to each slide then coverslipped. Sections were imaged on a confocal microscope (Zeiss 710) using a 20X objective.

### FISH quantification

Image quantification was performed using CellProfiler v4.0.4 (Stirling et al., 2021). DAPI-stained nuclei were first identified using the ClassifyPixels-Unet module. Areas corresponding to cells were estimated by extending nuclei boundaries by 5 pixels. Then signal puncta for each channel were identified and their intensities were measured. For each cell and each channel, we calculated the summed signal intensity of overlapping puncta divided by the cell area. To test for significant differences in gene expression between hearing and deaf birds, a linear mixed effects model was fit using function *lmer* from R package *lmerTest v3.1-3 (Kuznetsova et al., 2017)* for each target gene and brain region as ‘intensity ~ condition + (1|bird)’ where ‘condition’ is contra or ipsi and ‘(1|bird)’ is the per-bird grouping factor. P-values were obtained by comparing this model with a reduced model ‘intensity ~ (1|bird)’ using ANOVA.

### Data availability

SLCR-seq mapped sequencing reads, gene-by-sample count matrices, and metadata can be found at NCBI GEO for deafening (accession number GSE200663) and unilateral LMAN lesion datasets (GSE200664).

### Code availability

Code underlying the analysis of birdsong and SLCR-seq gene expression can be found in the GitHub repository https://github.com/bradleycolquitt/deaf_gex.

## Acknowledgements

We would like to thank Andrea Hausenstaub and Christoph Schreiner for providing critical commentary on this manuscript, Adria Arteseros for providing technical expertise, and Mimi Kao for surgical expertise.

## Supplemental Tables

***Supplemental Table 1. SLCR-seq barcode and index sequences.***

Full sequences for RT_primer_v1, RT_primer_v2, and PCR2.

***Supplemental Table 2. deafening voom statistics***

Fold-change estimates and p-values for voom regression analysis of the deafening SLCR-seq dataset

***Supplemental Table 3. GSEA statistics***

Gene set enrichment analysis of song destabilization-associated genes

***Supplemental Table 4. Network module memberships***

Gene memberships in MEGENA modules for the RA, HVC, LMAN, and Area X networks

***Supplemental Table 5. Differential correlation analysis***

Inter-region gene correlations results for the combined dataset and split between hearing and deaf birds

***Supplemental Table 6. Unilateral LMAN lesion voom statistics***

Fold-change estimates and p-values for voom regression analysis of the unilateral LMAN lesion SLCR-seq dataset

## References

Aldenhoff JB, Gruol DL, Rivier J, Vale W, Siggins GR. 1983. Corticotropin releasing factor decreases postburst hyperpolarizations and excites hippocampal neurons. Science 221:875–877. doi:10.1126/science.6603658

Andalman AS, Fee MS. 2009. A basal ganglia-forebrain circuit in the songbird biases motor output to avoid vocal errors. Proc Natl Acad Sci U S A 106:12518–12523. doi:10.1073/pnas.0903214106

Bargmann CI. 2012. Beyond the connectome: How neuromodulators shape neural circuits. Bioessays 34:458–465. doi:10.1002/bies.201100185

Bates D, Mächler M, Bolker B, Walker S. 2015. Fitting Linear Mixed-Effects Models Using lme4. J Stat Softw 67:1–48. doi:10.18637/jss.v067.i01

Blank T, Nijholt I, Grammatopoulos DK, Randeva HS, Hillhouse EW, Spiess J. 2003. Corticotropin-releasing factor receptors couple to multiple G-proteins to activate diverse intracellular signaling pathways in mouse hippocampus: role in neuronal excitability and associative learning. J Neurosci 23:700–707. doi:10.1523/JNEUROSCI.23-02-00700.2003

Bottjer SW, Miesner EA, Arnold AP. 1984. Forebrain lesions disrupt development but not maintenance of song in passerine birds. Science 224:901–903. doi:10.1126/science.6719123

Brainard MS, Doupe AJ. 2013. Translating birdsong: songbirds as a model for basic and applied medical research. Annu Rev Neurosci 36:489–517.

Brainard MS, Doupe AJ. 2001. Postlearning consolidation of birdsong: stabilizing effects of age and anterior forebrain lesions. J Neurosci 21:2501–17.

Brainard MS, Doupe AJ. 2000. Interruption of a basal ganglia-forebrain circuit prevents plasticity of learned vocalizations. Nature 404:762–766. doi:10.1038/35008083

Bray NL, Pimentel H, Melsted P, Pachter L. 2016. Near-optimal probabilistic RNA-seq quantification. Nat Biotechnol 34:525–527. doi:10.1038/nbt.3519

Charlesworth JD, Warren TL, Brainard MS. 2012. Covert skill learning in a cortical-basal ganglia circuit. Nature 486:251–255.

Chen DY, Stern SA, Garcia-Osta A, Saunier-Rebori B, Pollonini G, Bambah-Mukku D, Blitzer RD, Alberini CM. 2011. A critical role for IGF-II in memory consolidation and enhancement. Nature 469:491–497. doi:10.1038/nature09667

Chen Q, Heston JB, Burkett ZD, White SA. 2013. Expression analysis of the speech-related genes FoxP1 and FoxP2 and their relation to singing behavior in two songbird species. J Exp Biol 216:3682–3692. doi:10.1242/jeb.085886

Chen SX, Kim AN, Peters AJ, Komiyama T. 2015. Subtype-specific plasticity of inhibitory circuits in motor cortex during motor learning. Nat Neurosci 18:1109–15. doi:10.1038/nn.4049

Chen Y, Dubé CM, Rice CJ, Baram TZ. 2008. Rapid loss of dendritic spines after stress involves derangement of spine dynamics by corticotropin-releasing hormone. J Neurosci 28:2903–2911. doi:10.1523/JNEUROSCI.0225-08.2008

Chu Y, Fioravante D, Leitges M, Regehr WG. 2014. Calcium-dependent PKC isoforms have specialized roles in short-term synaptic plasticity. Neuron 82:859–71. doi:10.1016/j.neuron.2014.04.003

Colquitt BM, Merullo DP, Konopka G, Roberts TF, Brainard MS. 2021. Cellular transcriptomics reveals evolutionary identities of songbird vocal circuits. Science 371. doi:10.1126/science.abd9704

Cowie R, Douglas-Cowie E, Kerr AG. 1982. A study of speech deterioration in post-lingually deafened adults. J Laryngol Otol 96:101–112. doi:10.1017/s002221510009229x

Curmi PA, Andersen SS, Lachkar S, Gavet O, Karsenti E, Knossow M, Sobel A. 1997. The stathmin/tubulin interaction in vitro. J Biol Chem 272:25029–25036. doi:10.1074/jbc.272.40.25029

Dobin A, Davis CA, Schlesinger F, Drenkow J, Zaleski C, Jha S, Batut P, Chaisson M, Gingeras TR. 2013. STAR: ultrafast universal RNA-seq aligner. Bioinformatics 29:15–21.

Donato F, Rompani SB, Caroni P. 2013. Parvalbumin-expressing basket-cell network plasticity induced by experience regulates adult learning. Nature 504:272–276. doi:10.1038/nature12866

Feenders G, Liedvogel M, Rivas M, Zapka M, Horita H, Hara E, Wada K, Mouritsen H, Jarvis ED. 2008. Molecular Mapping of Movement-Associated Areas in the Avian Brain: A Motor Theory for Vocal Learning Origin. PLoS One 3:e1768. doi:10.1371/journal.pone.0001768

Finak G, McDavid A, Yajima M, Deng J, Gersuk V, Shalek AK, Slichter CK, Miller HW, McElrath MJ, Prlic M, Linsley PS, Gottardo R. 2015. MAST: a flexible statistical framework for assessing transcriptional changes and characterizing heterogeneity in single-cell RNA sequencing data. Genome Biol 16:278. doi:10.1186/s13059-015-0844-5

Fioravante D, Chu Y, de Jong AP, Leitges M, Kaeser PS, Regehr WG. 2014. Protein kinase C is a calcium sensor for presynaptic short-term plasticity. Elife 3:e03011. doi:10.7554/eLife.03011

Fox EA, Gruol DL. 1993. Corticotropin-releasing factor suppresses the afterhyperpolarization in cerebellar Purkinje neurons. Neurosci Lett 149:103–107. doi:10.1016/0304-3940(93)90358-r

Fox J, Weisberg S. 2019. An R Companion to Applied Regression.

Fukushima M, Margoliash D. 2015. The effects of delayed auditory feedback revealed by bone conduction microphone in adult zebra finches. Sci Rep 5:8800. doi:10.1038/srep08800

Garst-Orozco J, Babadi B, Ölveczky BP. 2014. A neural circuit mechanism for regulating vocal variability during song learning in zebra finches. Elife 3:e03697. doi:10.7554/eLife.03697

Gilliam DT, Menon V, Bretz NP, Pruszak J. 2017. The CD24 surface antigen in neural development and disease. Neurobiol Dis 99:133–144. doi:10.1016/j.nbd.2016.12.011

Goldberg JH, Fee MS. 2012. A cortical motor nucleus drives the basal ganglia-recipient thalamus in singing birds. Nat Neurosci 15:620–7. doi:10.1038/nn.3047

Hamaguchi K, Tschida KA, Yoon I, Donald BR, Mooney R. 2014. Auditory synapses to song premotor neurons are gated off during vocalization in zebra finches. Elife 3:e01833. doi:10.7554/eLife.01833

Hayase S, Wang H, Ohgushi E, Kobayashi M, Mori C, Horita H, Mineta K, Liu W-C, Wada K. 2018. Vocal practice regulates singing activity-dependent genes underlying age-independent vocal learning in songbirds. PLoS Biol 16:e2006537. doi:10.1371/journal.pbio.2006537

Helduser S, Cheng S, Güntürkün O. 2013. Identification of two forebrain structures that mediate execution of memorized sequences in the pigeon. J Neurophysiol 109:958–968. doi:10.1152/jn.00763.2012

Hilliard AT, Miller JE, Fraley ER, Horvath S, White SA. 2012. Molecular microcircuitry underlies functional specification in a basal ganglia circuit dedicated to vocal learning. Neuron 73:537–552. doi:10.1016/j.neuron.2012.01.005

Horita H, Kobayashi M, Liu W-C, Oka K, Jarvis ED, Wada K. 2012. Specialized motor-driven dusp1 expression in the song systems of multiple lineages of vocal learning birds. PLoS One 7:e42173. doi:10.1371/journal.pone.0042173

Hou Z-H, Yu X. 2013. Activity-regulated somatostatin expression reduces dendritic spine density and lowers excitatory synaptic transmission via postsynaptic somatostatin receptor 4. J Biol Chem 288:2501–2509. doi:10.1074/jbc.M112.419051

Islam S, Kjällquist U, Moliner A, Zajac P, Fan J-B, Lönnerberg P, Linnarsson S. 2012. Highly multiplexed and strand-specific single-cell RNA 5’ end sequencing. Nat Protoc 7:813–28. doi:10.1038/nprot.2012.022

Islam S, Zeisel A, Joost S, La Manno G, Zajac P, Kasper M, Lönnerberg P, Linnarsson S. 2014. Quantitative single-cell RNA-seq with unique molecular identifiers. Nat Methods 11:163–6. doi:10.1038/nmeth.2772

Jarvis ED, Scharff C, Grossman MR, Ramos JA, Nottebohm F. 1998. For whom the bird sings: context-dependent gene expression. Neuron 21:775–788. doi:10.1016/s0896-6273(00)80594-2

Jevsek M, Jaworski A, Polo-Parada L, Kim N, Fan J, Landmesser LT, Burden SJ. 2006. CD24 is expressed by myofiber synaptic nuclei and regulates synaptic transmission. Proc Natl Acad Sci U S A 103:6374–6379. doi:10.1073/pnas.0601468103

Johnson WE, Li C, Rabinovic A. 2007. Adjusting batch effects in microarray expression data using empirical Bayes methods. Biostatistics 8:118–27. doi:10.1093/biostatistics/kxj037

Kao MH, Brainard MS. 2006. Lesions of an avian basal ganglia circuit prevent context-dependent changes to song variability. J Neurophysiol 96:1441–1455.

Kao MH, Doupe AJ, Brainard MS. 2005. Contributions of an avian basal ganglia-forebrain circuit to real-time modulation of song. Nature 433:638–643.

Kemp CF, Woods RJ, Lowry PJ. 1998. The corticotrophin-releasing factor-binding protein: an act of several parts. Peptides 19:1119–1128. doi:10.1016/s0196-9781(98)00057-6

Kivioja T, Vähärautio A, Karlsson K, Bonke M, Enge M, Linnarsson S, Taipale J. 2012. Counting absolute numbers of molecules using unique molecular identifiers. Nat Methods 9:72–4. doi:10.1038/nmeth.1778

Kojima S, Kao MH, Doupe AJ. 2013. Task-related “cortical” bursting depends critically on basal ganglia input and is linked to vocal plasticity. Proc Natl Acad Sci U S A 110:4756–4761. doi:10.1073/pnas.1216308110

Korotkevich G, Sukhov V, Budin N, Shpak B, Artyomov MN, Sergushichev A. 2021. Fast gene set enrichment analysis. bioRxiv. doi:10.1101/060012

Kosche G, Vallentin D, Long MA. 2015. Interplay of inhibition and excitation shapes a premotor neural sequence. J Neurosci 35:1217–1227. doi:10.1523/JNEUROSCI.4346-14.2015

Kotak VC, Fujisawa S, Lee FA, Karthikeyan O, Aoki C, Sanes DH. 2005. Hearing loss raises excitability in the auditory cortex. J Neurosci 25:3908–3918. doi:10.1523/JNEUROSCI.5169-04.2005

Kotak VC, Takesian AE, Sanes DH. 2008. Hearing loss prevents the maturation of GABAergic transmission in the auditory cortex. Cereb Cortex 18:2098–2108. doi:10.1093/cercor/bhm233

Kovaka S, Zimin AV, Pertea GM, Razaghi R, Salzberg SL, Pertea M. 2019. Transcriptome assembly from long-read RNA-seq alignments with StringTie2. Genome Biol 20:278. doi:10.1186/s13059-019-1910-1

Kratzer S, Mattusch C, Metzger MW, Dedic N, Noll-Hussong M, Kafitz KW, Eder M, Deussing JM, Holsboer F, Kochs E, Rammes G. 2013. Activation of CRH receptor type 1 expressed on glutamatergic neurons increases excitability of CA1 pyramidal neurons by the modulation of voltage-gated ion channels. Front Cell Neurosci 7:91. doi:10.3389/fncel.2013.00091

Kröner S, Güntürkün O. 1999. Afferent and efferent connections of the caudolateral neostriatum in the pigeon (Columba livia): A retro- and anterograde pathway tracing study. J Comp Neurol 407:228–260. doi:10.1002/(SICI)1096-9861(19990503)407:2<228::AID-CNE6>3.0.CO;2-2

Kuznetsova A, Brockhoff PB, Christensen RHB. 2017. lmerTest Package: Tests in Linear Mixed Effects Models. Journal of Statistical Software. doi:10.18637/jss.v082.i13

Lane H, Webster JW. 1991. Speech deterioration in postlingually deafened adults. J Acoust Soc Am 89:859–866. doi:10.1121/1.1894647

Langfelder P, Horvath S. 2008. WGCNA: an R package for weighted correlation network analysis. BMC Bioinformatics 9:559. doi:10.1186/1471-2105-9-559

Langfelder P, Luo R, Oldham MC, Horvath S. 2011. Is my network module preserved and reproducible? PLoS Comput Biol 7:e1001057. doi:10.1371/journal.pcbi.1001057

Law CW, Chen Y, Shi W, Smyth GK. 2014. voom: Precision weights unlock linear model analysis tools for RNA-seq read counts. Genome Biol 15:R29. doi:10.1186/gb-2014-15-2-r29

Leek JT. 2014. svaseq: removing batch effects and other unwanted noise from sequencing data. Nucleic Acids Res 42. doi:10.1093/nar/gku864

Leonardo A, Konishi M. 1999. Decrystallization of adult birdsong by perturbation of auditory feedback. Nature 399:466–70. doi:10.1038/20933

Li K, Nakajima M, Ibañez-Tallon I, Heintz N. 2016. A Cortical Circuit for Sexually Dimorphic Oxytocin-Dependent Anxiety Behaviors. Cell 167:60–72.e11. doi:10.1016/j.cell.2016.08.067

Livingston FS, White SA, Mooney R. 2000. Slow NMDA-EPSCs at synapses critical for song development are not required for song learning in zebra finches. Nat Neurosci 3:482–488. doi:10.1038/74857

Lombardino AJ, Nottebohm F. 2000. Age at deafening affects the stability of learned song in adult male zebra finches. J Neurosci 20:5054–5064. doi:10.1523/JNEUROSCI.20-13-05054.2000

Lovell PV, Wirthlin M, Kaser T, Buckner AA, Carleton JB, Snider BR, McHugh AK, Tolpygo A, Mitra PP, Mello CV. 2020. ZEBrA: Zebra finch Expression Brain Atlas-A resource for comparative molecular neuroanatomy and brain evolution studies. J Comp Neurol 528:2099–2131. doi:10.1002/cne.24879

Macosko EZ, Basu A, Satija R, Nemesh J, Shekhar K, Goldman M, Tirosh I, Bialas AR, Kamitaki N, Martersteck EM, Trombetta JJ, Weitz DA, Sanes JR, Shalek AK, Regev A, McCarroll SA. 2015. Highly Parallel Genome-wide Expression Profiling of Individual Cells Using Nanoliter Droplets. Cell 161:1202–1214. doi:10.1016/j.cell.2015.05.002

Mandelblat-Cerf Y, Las L, Denisenko N, Fee MS. 2014. A role for descending auditory cortical projections in songbird vocal learning. Elife 3. doi:10.7554/eLife.02152

Marder E. 2011. Variability, compensation, and modulation in neurons and circuits. Proc Natl Acad Sci U S A 108 Suppl:15542–8. doi:10.1073/pnas.1010674108

Melsted P, Sina Booeshaghi A, Liu L, Gao F, Lu L, Min KH (joseph), da Veiga Beltrame E, Hjörleifsson KE, Gehring J, Pachter L. 2021. Modular, efficient and constant-memory single-cell RNA-seq preprocessing. Nat Biotechnol 1–6. doi:10.1038/s41587-021-00870-2

Mets DG, Brainard MS. 2018. An automated approach to the quantitation of vocalizations and vocal learning in the songbird. PLoS Comput Biol 14:e1006437. doi:10.1371/journal.pcbi.1006437

Miller JE, Hilliard AT, White SA. 2010. Song practice promotes acute vocal variability at a key stage of sensorimotor learning. PLoS One 5:e8592. doi:10.1371/journal.pone.0008592

Miller MN, Cheung CYJ, Brainard MS. 2017. Vocal learning promotes patterned inhibitory connectivity. Nat Commun 8:2105. doi:10.1038/s41467-017-01914-5

Moorman S, Ahn J-R, Kao MH. 2021. Plasticity of stereotyped birdsong driven by chronic manipulation of cortical-basal ganglia activity. Curr Biol 31:2619–2632.e4. doi:10.1016/j.cub.2021.04.030

Nevue AA, Lovell PV, Wirthlin M, Mello CV. 2020. Molecular specializations of deep cortical layer analogs in songbirds. Sci Rep 10:18767. doi:10.1038/s41598-020-75773-4

Nicholson D. 2021. NickleDave/hybrid-vocal-classifier. doi:10.5281/zenodo.4678768

Nordeen KW, Nordeen EJ. 1993. Long-term maintenance of song in adult zebra finches is not affected by lesions of a forebrain region involved in song learning. Behav Neural Biol 59:79–82. doi:10.1016/0163-1047(93)91215-9

Nordeen KW, Nordeen EJ. 1992. Auditory feedback is necessary for the maintenance of stereotyped song in adult zebra finches. Behav Neural Biol 57:58–66. doi:10.1016/0163-1047(92)90757-U

Nottebohm F, Kelley DB, Paton JA. 1982. Connections of vocal control nuclei in the canary telencephalon. J Comp Neurol 207:344–57. doi:10.1002/cne.902070406

Nottebohm F, Stokes TM, Leonard CM. 1976. Central control of song in the canary, Serinus canarius. J Comp Neurol 165:457–86. doi:10.1002/cne.901650405

Ohgushi E, Mori C, Wada K. 2015. Diurnal oscillation of vocal development associated with clustered singing by juvenile songbirds. J Exp Biol 218:2260–2268. doi:10.1242/jeb.115105

Okanoya K, Yamaguchi A. 1997. Adult Bengalese finches (Lonchura striata var. domestica) require real-time auditory feedback to produce normal song syntax. J Neurobiol 33:343–56.

Olveczky BP, Andalman AS, Fee MS. 2005. Vocal experimentation in the juvenile songbird requires a basal ganglia circuit. PLoS Biol 3:e153. doi:10.1371/journal.pbio.0030153

Ölveczky BP, Otchy TM, Goldberg JH, Aronov D, Fee MS. 2011. Changes in the neural control of a complex motor sequence during learning. J Neurophysiol 106:386–397. doi:10.1152/jn.00018.2011

Peng Z, Zeng S, Liu Y, Dong Y, Zhang H, Zhang X, Zuo M. 2012a. Comparative study on song behavior, and ultra-structural, electrophysiological and immunoreactive properties in RA among deafened, untutored and normal-hearing Bengalese finches. Brain Res 1458:40–55. doi:10.1016/j.brainres.2012.04.009

Peng Z, Zhang X, Liu Y, Xi C, Zeng S, Zhang X, Zuo M, Xu J, Ji Y, Han Z. 2013. Ultrastructural and electrophysiological analysis of Area X in the untutored and deafened Bengalese finch in relation to normally reared birds. Brain Res 1527:87–98. doi:10.1016/j.brainres.2013.06.031

Peng Z, Zhang X, Xi C, Zeng S, Liu N, Zuo M, Zhang X. 2012b. Changes in ultra-structures and electrophysiological properties in HVC of untutored and deafened Bengalese finches relation to normally reared birds: implications for song learning. Brain Res Bull 89:211–222. doi:10.1016/j.brainresbull.2012.09.004

Pfenning AR, Hara E, Whitney O, Rivas MV, Wang R, Roulhac PL, Howard JT, Wirthlin M, Lovell PV, Ganapathy G, Mouncastle J, Moseley MA, Thompson JW, Soderblom EJ, Iriki A, Kato M, Gilbert MTP, Zhang G, Bakken T, Bongaarts A, Bernard A, Lein E, Mello CV, Hartemink AJ, Jarvis ED. 2014. Convergent transcriptional specializations in the brains of humans and song-learning birds. Science 346:1256846–1256846. doi:10.1126/science.1256846

Picelli S, Björklund \aasa K., Faridani OR, Sagasser S, Winberg G, Sandberg R. 2013. Smart-seq2 for sensitive full-length transcriptome profiling in single cells. Nat Methods 10:1096–8. doi:10.1038/nmeth.2639

Picelli S, Björklund AK, Reinius B, Sagasser S, Winberg G, Sandberg R. 2014. Tn5 transposase and tagmentation procedures for massively scaled sequencing projects. Genome Res 24:2033–40. doi:10.1101/gr.177881.114

Pittman QJ, Siggins GR. 1981. Somatostatin hyperpolarizes hippocampal pyramidal cells in vitro. Brain Res 221:402–408. doi:10.1016/0006-8993(81)90791-5

Pytte CL, George S, Korman S, David E, Bogdan D, Kirn JR. 2012. Adult neurogenesis is associated with the maintenance of a stereotyped, learned motor behavior. - PubMed - NCBI. doi:10.1523/JNEUROSCI.5385-11.2012

Ritchie ME, Phipson B, Wu D, Hu Y, Law CW, Shi W, Smyth GK. 2015. limma powers differential expression analyses for RNA-sequencing and microarray studies. Nucleic Acids Res 43:e47. doi:10.1093/nar/gkv007

Roy A, Mooney R. 2007. Auditory plasticity in a basal ganglia-forebrain pathway during decrystallization of adult birdsong. J Neurosci 27:6374–6387. doi:10.1523/JNEUROSCI.0894-07.2007

Sainburg T. 2020. Bengalese finch. doi:10.5281/zenodo.3926440

Sainburg T, Thielk M, Gentner TQ. 2020. Finding, visualizing, and quantifying latent structure across diverse animal vocal repertoires. PLoS Comput Biol 16:e1008228. doi:10.1371/journal.pcbi.1008228

Sakata JT, Hampton CM, Brainard MS. 2008. Social modulation of sequence and syllable variability in adult birdsong. J Neurophysiol 99:1700–1711.

Sasaki A, Sotnikova TD, Gainetdinov RR, Jarvis ED. 2006. Social context-dependent singing-regulated dopamine. J Neurosci 26:9010–9014.

Scharff C, Nottebohm F. 1991. A comparative study of the behavioral deficits following lesions of various parts of the zebra finch song system: implications for vocal learning. J Neurosci 11:2896–2913.

Schenk BS, Baumgartner WD, Hamzavi JS. 2003. Effect of the loss of auditory feedback on segmental parameters of vowels of postlingually deafened speakers. Auris Nasus Larynx 30:333–339. doi:10.1016/s0385-8146(03)00093-2

Schindelin J, Arganda-Carreras I, Frise E, Kaynig V, Longair M, Pietzsch T, Preibisch S, Rueden C, Saalfeld S, Schmid B, Tinevez J-Y, White DJ, Hartenstein V, Eliceiri K, Tomancak P, Cardona A. 2012. Fiji: an open-source platform for biological-image analysis. Nat Methods 9:676–682. doi:10.1038/nmeth.2019

Scott LL, Nordeen EJ, Nordeen KW. 2000. The relationship between rates of HVc neuron addition and vocal plasticity in adult songbirds. J Neurobiol 43:79–88. doi:10.1002/(sici)1097-4695(200004)43:1<79::aid-neu7>3.0.co;2-p

Scrucca L, Fop M, Murphy TB, Raftery AE. 2016. mclust 5: Clustering, Classification and Density Estimation Using Gaussian Finite Mixture Models. R J 8:289–317. doi:10.32614/RJ-2016-021

Seki S, Eggermont JJ. 2003. Changes in spontaneous firing rate and neural synchrony in cat primary auditory cortex after localized tone-induced hearing loss. Hear Res 180:28–38. doi:10.1016/s0378-5955(03)00074-1

Shumyatsky GP, Malleret G, Shin R-M, Takizawa S, Tully K, Tsvetkov E, Zakharenko SS, Joseph J, Vronskaya S, Yin D, Schubart UK, Kandel ER, Bolshakov VY. 2005. stathmin, a gene enriched in the amygdala, controls both learned and innate fear. Cell 123:697–709. doi:10.1016/j.cell.2005.08.038

Simpson HB, Vicario DS. 1990. Brain pathways for learned and unlearned vocalizations differ in zebra finches. J Neurosci 10:1541–1556.

Sohrabji F, Nordeen EJ, Nordeen KW. 1990. Selective impairment of song learning following lesions of a forebrain nucleus in the juvenile zebra finch. Behav Neural Biol 53:51–63. doi:10.1016/0163-1047(90)90797-a

Song W-M, Zhang B. 2015. Multiscale Embedded Gene Co-expression Network Analysis. PLoS Comput Biol 11:e1004574. doi:10.1371/journal.pcbi.1004574

Song Y-H, Yoon J, Lee S-H. 2021. The role of neuropeptide somatostatin in the brain and its application in treating neurological disorders. Exp Mol Med 53:328–338. doi:10.1038/s12276-021-00580-4

Stirling DR, Swain-Bowden MJ, Lucas AM, Carpenter AE, Cimini BA, Goodman A. 2021. CellProfiler 4: improvements in speed, utility and usability. BMC Bioinformatics 22:433. doi:10.1186/s12859-021-04344-9

Stuart T, Butler A, Hoffman P, Hafemeister C, Papalexi E, Mauck WM, Hao Y, Stoeckius M, Smibert P, Satija R. 2019. Comprehensive Integration of Single-Cell Data. Cell 177:1888–1902.e21. doi:10.1016/j.cell.2019.05.031

Sueur J, Aubin T, Simonis C. 2008. Seewave: a free modular tool for sound analysis and synthesis. Bioacoustics.

Tachibana RO, Oosugi N, Okanoya K. 2014. Semi-automatic classification of birdsong elements using a linear support vector machine. PLoS One 9:e92584. doi:10.1371/journal.pone.0092584

Todorov E. 2004. Optimality principles in sensorimotor control. Nat Neurosci 7:907–915. doi:10.1038/nn1309

Tschida KA, Mooney R. 2012. Deafening drives cell-type-specific changes to dendritic spines in a sensorimotor nucleus important to learned vocalizations. Neuron 73:1028–39. doi:10.1016/j.neuron.2011.12.038

Tyssowski KM, DeStefino NR, Cho J-H, Dunn CJ, Poston RG, Carty CE, Jones RD, Chang SM, Romeo P, Wurzelmann MK, Ward JM, Andermann ML, Saha RN, Dudek SM, Gray JM. 2018. Different Neuronal Activity Patterns Induce Different Gene Expression Programs. Neuron 98:530–546.e11. doi:10.1016/j.neuron.2018.04.001

Uchida S, Martel G, Pavlowsky A, Takizawa S, Hevi C, Watanabe Y, Kandel ER, Alarcon JM, Shumyatsky GP. 2014. Learning-induced and stathmin-dependent changes in microtubule stability are critical for memory and disrupted in ageing. Nat Commun 5:4389. doi:10.1038/ncomms5389

Vallentin D, Kosche G, Lipkind D, Long MA. 2016. Inhibition protects acquired song segments during vocal learning in zebra finches. Science 351:267–271. doi:10.1126/science.aad3023

Vicario DS. 1991. Organization of the zebra finch song control system: II. Functional organization of outputs from nucleus Robustus archistriatalis. J Comp Neurol 309:486–494. doi:10.1002/cne.903090405

Wada K, Howard JT, McConnell P, Whitney O, Lints T, Rivas MV, Horita H, Patterson MA, White SA, Scharff C, Haesler S, Zhao S, Sakaguchi H, Hagiwara M, Shiraki T, Hirozane-Kishikawa T, Skene P, Hayashizaki Y, Carninci P, Jarvis ED. 2006. A molecular neuroethological approach for identifying and characterizing a cascade of behaviorally regulated genes. Proc Natl Acad Sci U S A 103:15212–15217.

Waldstein RS. 1990. Effects of postlingual deafness on speech production: implications for the role of auditory feedback. J Acoust Soc Am 88:2099–2114. doi:10.1121/1.400107

Wang H, Eckel RH. 2012. Lipoprotein lipase in the brain and nervous system. Annu Rev Nutr 32:147–160. doi:10.1146/annurev-nutr-071811-150703

Wang N, Aviram R, Kirn JR. 1999. Deafening Alters Neuron Turnover within the Telencephalic Motor Pathway for Song Control in Adult Zebra Finches. J Neurosci 19:10554–10561. doi:10.1523/JNEUROSCI.19-23-10554.1999

Warren TL, Tumer EC, Charlesworth JD, Brainard MS. 2011. Mechanisms and time course of vocal learning and consolidation in the adult songbird. J Neurophysiol 106:1806–1821.

Warren WC, Clayton DF, Ellegren H, Arnold AP, Hillier LW, Künstner A, Searle S, White S, Vilella AJ, Fairley S, Heger A, Kong L, Ponting CP, Jarvis ED, Mello CV, Minx P, Lovell P, Velho TAF, Ferris M, Balakrishnan CN, Sinha S, Blatti C, London SE, Li Y, Lin Y-C, George J, Sweedler J, Southey B, Gunaratne P, Watson M, Nam K, Backström N, Smeds L, Nabholz B, Itoh Y, Whitney O, Pfenning AR, Howard J, Völker M, Skinner BM, Griffin DK, Ye L, McLaren WM, Flicek P, Quesada V, Velasco G, Lopez-Otin C, Puente XS, Olender T, Lancet D, Smit AFA, Hubley R, Konkel MK, Walker JA, Batzer MA, Gu W, Pollock DD, Chen L, Cheng Z, Eichler EE, Stapley J, Slate J, Ekblom R, Birkhead T, Burke T, Burt D, Scharff C, Adam I, Richard H, Sultan M, Soldatov A, Lehrach H, Edwards SV, Yang S-P, Li X, Graves T, Fulton L, Nelson J, Chinwalla A, Hou S, Mardis ER, Wilson RK. 2010. The genome of a songbird. Nature 464:757–762. doi:10.1038/nature08819

Watanabe A, Kimura T, Sakaguchi H. 2002. Expression of protein kinase C in song control nuclei of deafened adult male Bengalese finches. Neuroreport 13:127–32.

Watanabe A, Li R, Kimura T, Sakaguchi H. 2006. Lesions of an avian forebrain nucleus prevent changes in protein kinase C levels associated with deafening-induced vocal plasticity in adult songbirds. Eur J Neurosci 23:2447–2457. doi:10.1111/j.1460-9568.2006.04763.x

Whitney O, Johnson F. 2005. Motor-induced transcription but sensory-regulated translation of ZENK in socially interactive songbirds. J Neurobiol 65:251–9. doi:10.1002/neu.20187

Whitney O, Pfenning AR, Howard JT, Blatti CA, Liu F, Ward JM, Wang R, Audet J-N, Kellis M, Mukherjee S, Sinha S, Hartemink AJ, West AE, Jarvis ED. 2014. Core and region-enriched networks of behaviorally regulated genes and the singing genome. Science 346:1256780–1256780. doi:10.1126/science.1256780

Wild JM. 1993. Descending projections of the songbird nucleus robustus archistriatalis. J Comp Neurol 338:225–241. doi:10.1002/cne.903380207

Williams H, Mehta N. 1999. Changes in adult zebra finch song require a forebrain nucleus that is not necessary for song production. J Neurobiol 39:14–28. doi:10.1002/(SICI)1097-4695(199904)39:1<14::AID-NEU2>3.0.CO;2-X

Wood JM, Lacherez P, Black AA, Cole MH, Boon MY, Kerr GK. 2011. Risk of falls, injurious falls, and other injuries resulting from visual impairment among older adults with age-related macular degeneration. Invest Ophthalmol Vis Sci 52:5088–5092. doi:10.1167/iovs.10-6644

Woolley SM, Rubel EW. 1997. Bengalese finches Lonchura Striata domestica depend upon auditory feedback for the maintenance of adult song. J Neurosci 17:6380–90.

Xian X, Liu T, Yu J, Wang Y, Miao Y, Zhang J, Yu Y, Ross C, Karasinska JM, Hayden MR, Liu G, Chui D. 2009. Presynaptic defects underlying impaired learning and memory function in lipoprotein lipase-deficient mice. J Neurosci 29:4681–4685. doi:10.1523/JNEUROSCI.0297-09.2009

Yu T, Taussig MD, DiPatrizio NV, Astarita G, Piomelli D, Bergman BC, Dell’Acqua ML, Eckel RH, Wang H. 2015. Deficiency of Lipoprotein Lipase in Neurons Decreases AMPA Receptor Phosphorylation and Leads to Neurobehavioral Abnormalities in Mice. PLoS One 10:e0135113. doi:10.1371/journal.pone.0135113

Zhou X, Fu X, Lin C, Zhou X, Liu J, Wang L, Zhang X, Zuo M, Fan X, Li D, Sun Y. 2017. Remodeling of Dendritic Spines in the Avian Vocal Motor Cortex Following Deafening Depends on the Basal Ganglia Circuit. Cereb Cortex 27:2820–2830. doi:10.1093/cercor/bhw130

